# Increased TGF β /Activin-Smad2 signaling is associated with pancreatic β -cell dysfunction and glucose intolerance in gestational diabetes mellitus

**DOI:** 10.1101/2025.02.16.638404

**Authors:** Talía Boronat-Belda, Hilda Ferrero, Sergi Soriano, Elena Ribes-García, Rubén Betoret-Gustems, Daniel Martínez-Bañón, Mónica Serrano-Selva, Juan Martínez-Pinna, Ángel Nadal, Iván Quesada, Paloma Alonso- Magdalena

## Abstract

**Background:** Gestational diabetes mellitus (GDM) is the most common metabolic disease during pregnancy and increases the prevalence of type 2 diabetes in both mothers and offspring. GDM management provides a window of opportunity to prevent and lower the global burden of diabetes across life. Molecular mechanisms underlying GDM are poorly defined. In this study, we explore the potential involvement of transforming growth factor beta (TGF-β) signaling in GDM as this pathway has been reported to affect pancreatic β-cell development, proliferation and identity.

**Methods:** We developed a GDM animal model. Serum circulating levels of TGFβ family ligands were measured in mice and human GDM. Pancreatic TGFβ signaling was investigated at the level of gene and protein expression.

**Results:** Our GDM animal model recapitulates the main pathophysiological features of human GDM including glucose intolerance, decreased insulin sensitivity and pancreatic β-cell malfunction. Islets from GDM mice showed impaired insulin secretion and content, altered ion channel activity, and decreased β-cell replication rate. This was accompanied by increased Smad2 signaling activation. Elevated serum activin-A and inhibin levels were found in mice and human GDM, suggesting their role as upstream signaling transducers of pancreatic Smad2 activation. Pharmacological inhibition of TGFβ/Activin-Smad2 signaling in mouse pancreatic islets resulted in improved pancreatic β-cell function and regeneration capacity of β-cells.

**Conclusions:** Our data disclose that disruption of pancreatic Smad2 pathway plays a critical role in the pathogenesis of GDM, contributing to abnormal glucose homeostasis and inadequate insulin secretion. Attenuation of this signaling pathway could represent a putative therapeutic target for GDM.

**Highlights:**

- High fat diet just before and during pregnancy leads to gestational diabetes in mice.
- Activin and inhibin serum levels are increased in human and mice gestational diabetes.
- Enhanced pancreatic Smad2 signaling contributes to inadequate insulin secretion in gestational diabetes.
- Inhibition of Smad2 signaling improves pancreatic β-cell function and proliferation.

## 1. Introduction

Gestational diabetes mellitus (GDM) is defined as any degree of glucose intolerance with onset or first recognition during pregnancy [1]. It is considered one of the most common medical complications of the gestational period [2]. The global standardized prevalence of GDM has increased considerably over the last decades and currently stands at 14.2 % [3], although GDM incidence has been reported as high as 27.0 % in certain regions like South East Asia [3]. While in most cases normal glucose tolerance is restored after delivery, women with GDM are at increased risk of developing type 2 diabetes mellitus (T2DM) in the postpartum period, highlighting that GDM diagnosis also carries long-term metabolic implications [4]. Accordingly, long-term follow up studies show that up to 20-60 % of women with GDM will have diabetes within 5-10 years after giving birth, and approximately 10% will have it shortly after delivery [5]. Besides, women who develop GDM have a 10-fold higher risk of progressing to T2DM compared to their peers [6]. From this life course perspective, GDM can be considered a chronic maternal metabolic disorder. Moreover, evidence to date confirm that GDM also places the offspring at a higher risk of developing metabolic disorders including T2DM and obesity [7]. Therefore, the road from GDM to T2DM represents a well-recognized continuum, which provide a unique opportunity to prevent and lower long-term burden of metabolic disorders in affected individuals.

Despite the importance of GDM the molecular mechanisms underlying the disease are poorly understood. During normal pregnancy, there is a progressive decline in maternal insulin sensitivity. This is a physiological response that serves as a protective mechanism to ensure the adequate nutrients supply to the growing foetus. Maternal euglycemia is maintained as insulin resistance is counterbalanced by a compensatory response of pancreatic β-cell function and mass whose failure is manifested as GDM [8]. These adaptive responses are thought to be driven, at least in part, by increased production of maternal hormones such as 17β-estradiol, progesterone, human placental lactogen (hPL) and prolactin [8–10]. Potential causes of inadequate β-cell function and failure in GDM remain elusive [11].

The transforming growth factor-beta (TGF-β) family of cytokines, comprises a number of highly conserved and structurally related proteins which regulate a wide variety of important biological process including cell growth, differentiation, apoptosis, extracellular matrix remodelling and fibrosis [12]. Based on sequence similarity, TGF-β, bone morphogenic proteins (BMPs) and activin/inhibin are considered the three major subgroups while Nodal and related factors form a separate subgroup mainly implicated in mesoderm and endoderm specification [13]. All the members of the TGF-β superfamily signal through a serine/threonine kinase receptor system which involves type-I and type-II receptors. In brief, ligand binding induces activation of the serine/threonine kinase of RII domain, which then phosphorylates RI on specific serine and threonine residues. Then, the activated type I receptor phosphorylates receptor-regulated Smad proteins (R-Smads), Smad1, 5 and 8 for BMP signaling, and Smad2 and 3 for TGFβ/Activin signaling. Activated R-Smad binds to the co-Smad, Smad4, forming heteromeric complexes which translocate into the nucleus. In the nucleus the Smad complex regulates the transcription of target genes [14].

TGFβ ligands, their receptors and Smad proteins are present in the endocrine pancreas. It has been stablished that TGFβ superfamily signaling regulates certain aspects of pancreas development, pancreatic β-cell proliferation, apoptosis and function [15, 16]. However, an important number of questions remain regarding how TGFβ signaling influences normal adult β-cell function and glucose homeostasis. Emerging evidence indicate that attenuation of TGFβ signaling may have protective effects against diabetes and obesity [16–19]. Nevertheless, the role of TGFβ signaling in GDM has not yet been explored.

## 2. Material and methods

### 2.1 Experimental methods Animals

C57BL/6 female mice (9-10 weeks of age) were used. C57BL/6 mice were obtained from Envigo (C57BL/6JOlaHsd; Barcelona, Spain) for breeding by the animal experimentation service of our institution that supplied the animals for the study. Animals were kept under controlled and standardized conditions and ad libitum access to food and water. All protocols were evaluated and approved by the Animal Ethics Committee of our institution in accordance with current national and European legislation (approvals ID: UMH.PAM.IB.03.18 and UMH.IDI.PA.240118).

The same week-old female mice were randomized to the different experimental groups based on body weight. For GDM model, mice were fed standard control diet (10 % energy from fat; D12450J, Research Diet, USA) or high fat diet (HFD) (60% energy from fat; D12492, Research Diet, USA) for three days prior to mating and throughout pregnancy. Pregnancy was determined by the presence of a vaginal plug the next morning of mating, which was identified as gestational day (GD) 0. Experiments were performed at indicated time points in the figure legends.

Non-pregnant mice were fed control diet or HFD for the equivalent length of time.

For ex vivo experiments adult female mice C57BL/6 (same strain as for in vivo) (8-10 weeks old) were used.

Additional details of materials and methods are provided as Supplemental Materials and Methods.

### 2.2 Statistical Analysis

After testing for normality, we performed Student’s t-test, one-way analysis of variance, or 2-way analysis of variance as required, when variables were normally distributed, and Kruskal–Wallis test, when variables were not normally distributed. Statistical analysis was performed with GraphPad Prism Software 7.0 (Graphpad Software, USA). Data are shown as mean ± SEM, and statistical significance was set at P< 0.05 for all analyses. The statistical tests used in each experiment were indicated in the corresponding figure legend.

## 3. Results

### 3.1. Glucose homeostasis in a GDM animal model and human GDM patients

Female mice were fed with a control diet or HFD three days before mating and along pregnancy. These experimental groups will be referred to throughout the text as control pregnant mice (CT-P) or HFD-pregnant mice (HFD-P) respectively. In parallel, non-pregnant mice were fed with either control diet or HFD during the equivalent length of time to that of pregnant mice. These groups will be named CT-NP and HFD-NP respectively.

We first studied the impact of high fat feeding in whole-body glucose homeostasis. To this end, an intraperitoneal glucose tolerance test (IPGTT) was performed at gestational day 15 (GD15) in pregnant mice or equivalent time in non-pregnant mice. We found that HFD administration resulted in marked glucose intolerance in HFD-P mice compared to CT-P, CT-NP and HFD-NP (Fig. 1A). This impaired glucose tolerance was also manifested in a significantly higher glucose AUC over the course of the IPGTT (Fig. 1B). HFD-NP mice also exhibited some degree of glucose intolerance compared to CT-NP, although the effect was much more modest than that of HFD-P mice (Fig. 1A). As expected, CT-P mice showed slightly higher blood glucose levels in response to the glucose load compared to CT-NP [20]. We also observed that fasting glycaemia levels were higher in HFD mice compared to their respectively control groups (Fig. 1C). No changes on glucose tolerance were found prior to mating (Fig. S1A-B). Next, we measured insulin release in response to glucose administration as a surrogate of pancreatic β-cell function. For this purpose, the stimulation index was determined as the ratio of insulin levels at 10 or 30 minutes after glucose administration to levels at 0 minutes. We found that insulin secretion was markedly diminished in HFD-P compared to CT-P as well as to CT-NP and HFD-NP (Fig. 1D) at 10 minutes; the same tendency was observed at 30 minutes (Fig. 1D). Fasting serum insulin levels were higher in HFD-P compared to the other groups which is an indicator of insulin resistance (Fig. 1E). No significant differences were found in fed insulin levels beyond an increase associated to pregnancy (Fig. 1F). Similar results were found in C-peptide levels (Fig. S1C). To further explore insulin sensitivity, an intraperitoneal insulin tolerance test (IPITT) was performed. Insulin sensitivity was decreased in both HFD-NP and HFD-P mice (Fig. 1G-H). We also found that leptin serum levels were upregulated in pregnant mice although the effect was more pronounced in HFD-P (Fig. 1I). On the contrary adiponectin, which serves as insulin sensitizer, was clearly decreased in HFD-P compared to CT-P (Fig. 1J).

**Figure 1.**
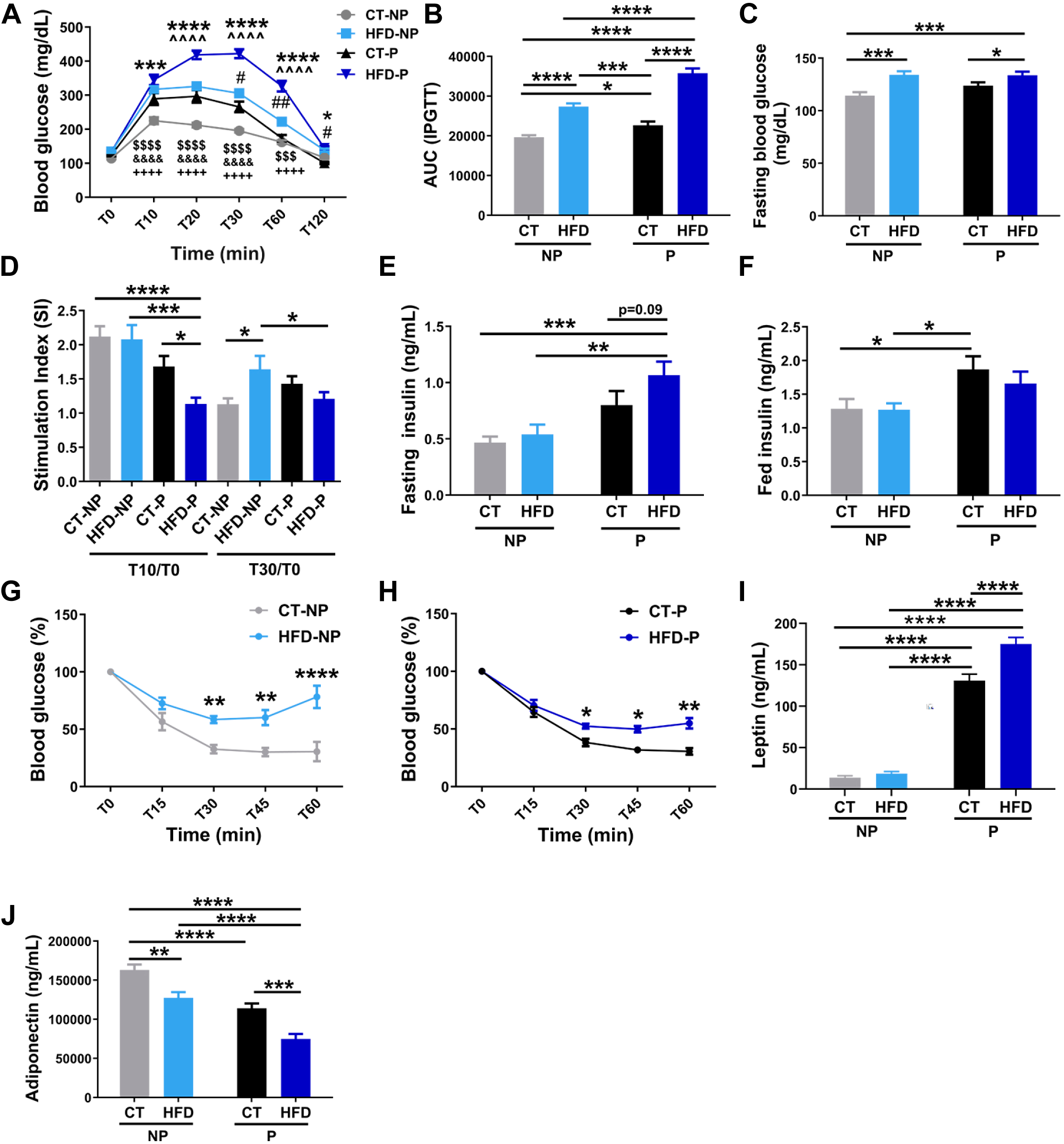
Study of glucose homeostasis and insulin sensitivity in control and GDM mice. **(A)** Intraperitoneal glucose tolerance test (IPGTT) in CT-NP (n=24), HFD-NP (n=23), CT-P (n=23) and HFD-P (n=23) mice at GD15 or equivalent length of time. **(B)** Area under the curve (AUC) from the IPGTT. **(C)** Fasting blood glucose levels in CT-NP (n=24), HFD-NP (n=23), CT-P (n=23) and HFD-P (n=23) mice. **(D)** In vivo insulin secretion in response to a glucose load determined as the ratio of insulin levels at 10 or 30 minutes to levels at 0 minutes for each mouse (CT-NP, n=18; HFD-NP, n=17; CT-P, n=14; HFD-P, n=26). **(E)** Fasting and **(F)** fed serum insulin levels (for fasting insulin: CT-NP, n=18; HFD-NP, n=16; CT-P, n=13; HFD-P, n=26; for fed insulin measurement: CT-NP, n=24; HFD-NP, n=21; CT-P, n=21; HFD-P, n=25). Intraperitoneal insulin tolerance test (IPITT) **(G)** in non-pregnant mice (CT-NP, n=6; HFD-NP, n=6) and **(H)** in pregnant-mice (CT-P, n=7; HFD-P, n=12) at GD15 or equivalent time period. **(I)** Leptin and **(J)** adiponectin serum levels (for leptin: CT-NP, n=16; HFD-NP, n=17; CT-P, n=18; HFD-P, n=18; for adiponectin: CT-NP, n=17; HFD-NP, n=17; CT-P, n=17; HFD-P, n=17). Data are presented as means ± SEM. Statistical comparisons were performed using Two-way ANOVA followed by Tukey’s (A, B, C, D, E, I, J), Fisher’s LSD (F) or Bonferroni’s (G, H) post hoc test. In (A) $$$ P < 0.001, $$$$ P < 0.0001 (CT-NP vs. HFD-NP); &&&& P < 0.0001 (CT-NP vs. CT-P); ++++ P < 0.0001 (CT-NP vs. HFD-P); *P < 0.05, ***P < 0.001, ****P < 0.0001 (CT-P vs. HFD-P); #P < 0.05, ##P < 0.01 (CT-P vs. HFD-NP); ^^^^ P < 0.0001 (HFD-NP vs. HFD-P). In the rest of panels *P < 0.05, **P < 0.01, ***P < 0.001 and ****P < 0.0001.

No changes on liver (Fig. S1D), mesenteric adipose (Fig. S1I) or pancreas (Fig. S1K) weight tissue were found in HFD-P. In contrast, increased weight of perigonadal (Fig. S1G) and decreased perirenal adipose tissue (Fig. S1J) was found in HFD-P compared to CT-P. Decreased muscle weight tissue (Fig. 1SE-F) and increased retroperitoneal weight tissue in HFD-P vs. non-pregnant mice was also found (Fig. S1H).

In a similar manner, we found that pregnant women diagnosed with GDM showed higher fasting insulin and C-peptide levels compared to healthy pregnant controls which is a strong indicator of insulin resistance (Fig 2A-B). In line with this, we found that HOMA-IR value was greater in GDM vs. control group (Fig. 2C). Triglyceride levels were also upregulated in GDM condition (Fig. 2D). Adiponectin, was significantly decreased in GDM (Fig. 2E). Moreover, Tumor necrosis factor-α (TNF-α) and Interleukin-6 (IL-6), pro-inflammatory cytokines associated to insulin resistance and diabetes, were increased in GDM group compared to controls (Fig. 2F-G).

**Figure 2.**
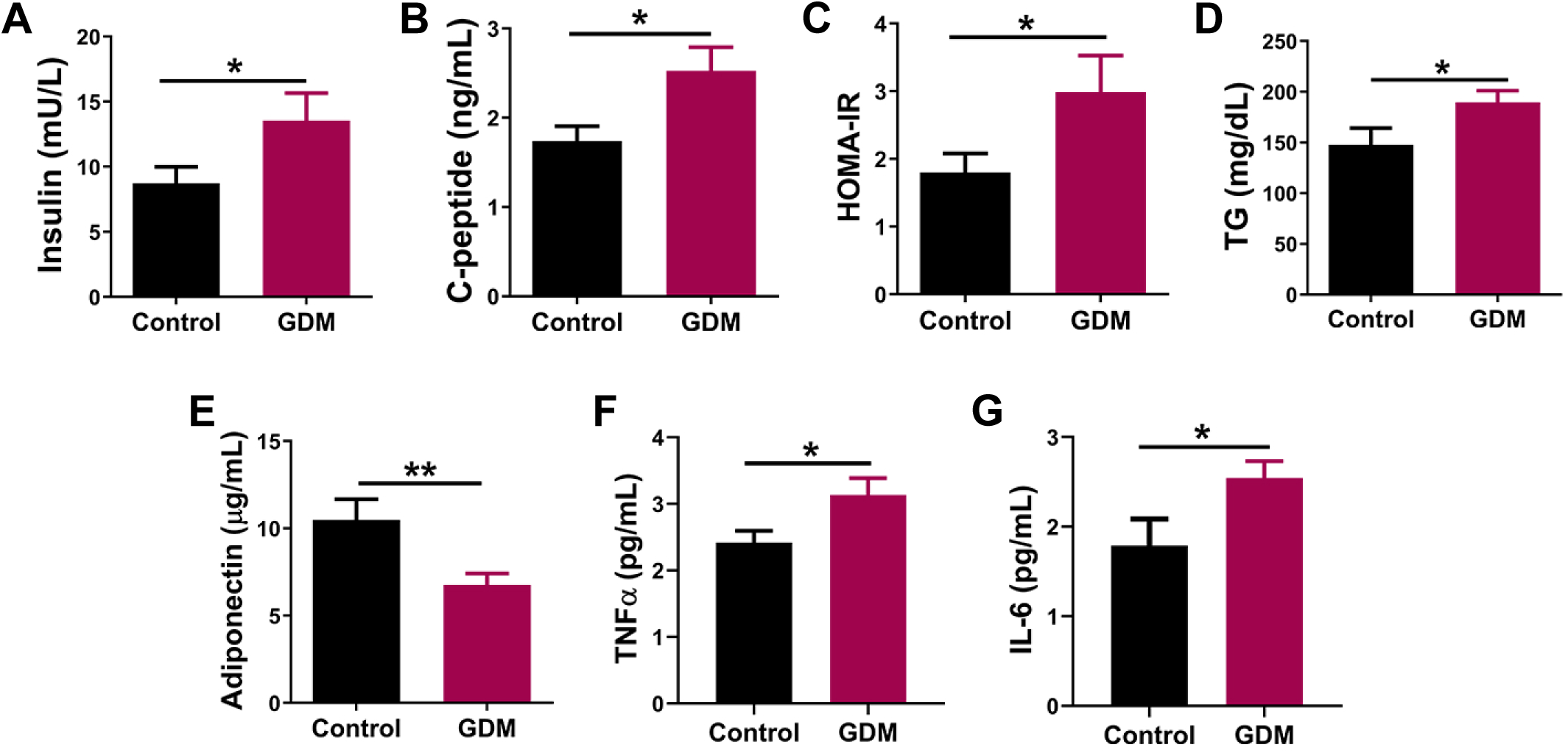
Serum measurements in Control and GDM patients. Serum fasting levels of **(A)** insulin and **(B)** C-peptide in control and GDM patients. **(C)** Quantification of HOMA-IR index. Serum levels of **(D)** TG, **(E)** adiponectin, **(F)** TNFα and **(G)** IL-6. Control (n=10-11), GDM (n= 13-14). Data are presented as means ± SEM. Statistical comparisons were performed using Student’s t test. *P < 0.05, **P < 0.01.

These findings indicate that HFD-P mice developed glucose intolerance, aggravated insulin resistance, altered cytokine levels and decreased pancreatic β-cell function, consistent with our findings in GDM patients and what is described in the literature [21–23]. These results indicate that this animal model is suitable for the study of GDM.

### 3.2. Study of pancreatic β-cell function and mass

Gene expression analysis was conducted in islets isolated from CT-NP, HFD-NP, CT-P and HFD-P mice with focus on genes relevant for function and identity of pancreatic β-cells. The most remarkable changes were decreased expression of *insulin* and *Mafa* genes in HFD-P compared to CT-P (Fig. 3A-B). Of note, *Mafa* expression was clearly upregulated in CT-P islets versus islets from non-pregnant mice (Fig. 3B). Furthermore, we found increased expression levels of *Pdx1* and *Hnf4a* (Fig. 3C-D) in pregnancy compared to non-pregnancy condition, the difference was even greater in HFD-P vs CT-P for *Hnf4a*. *Gck* gene expression was increased in HFD-P compared to HFD-NP (Fig. 3E) while no differences were found in *Glut2* gene (Fig. 3F). *Kir6.2* and *Sur1* gene expression was upregulated in islets isolated from HFD-P mice compared to non-pregnant islets (Fig. 3G-H).

**Figure 3.**
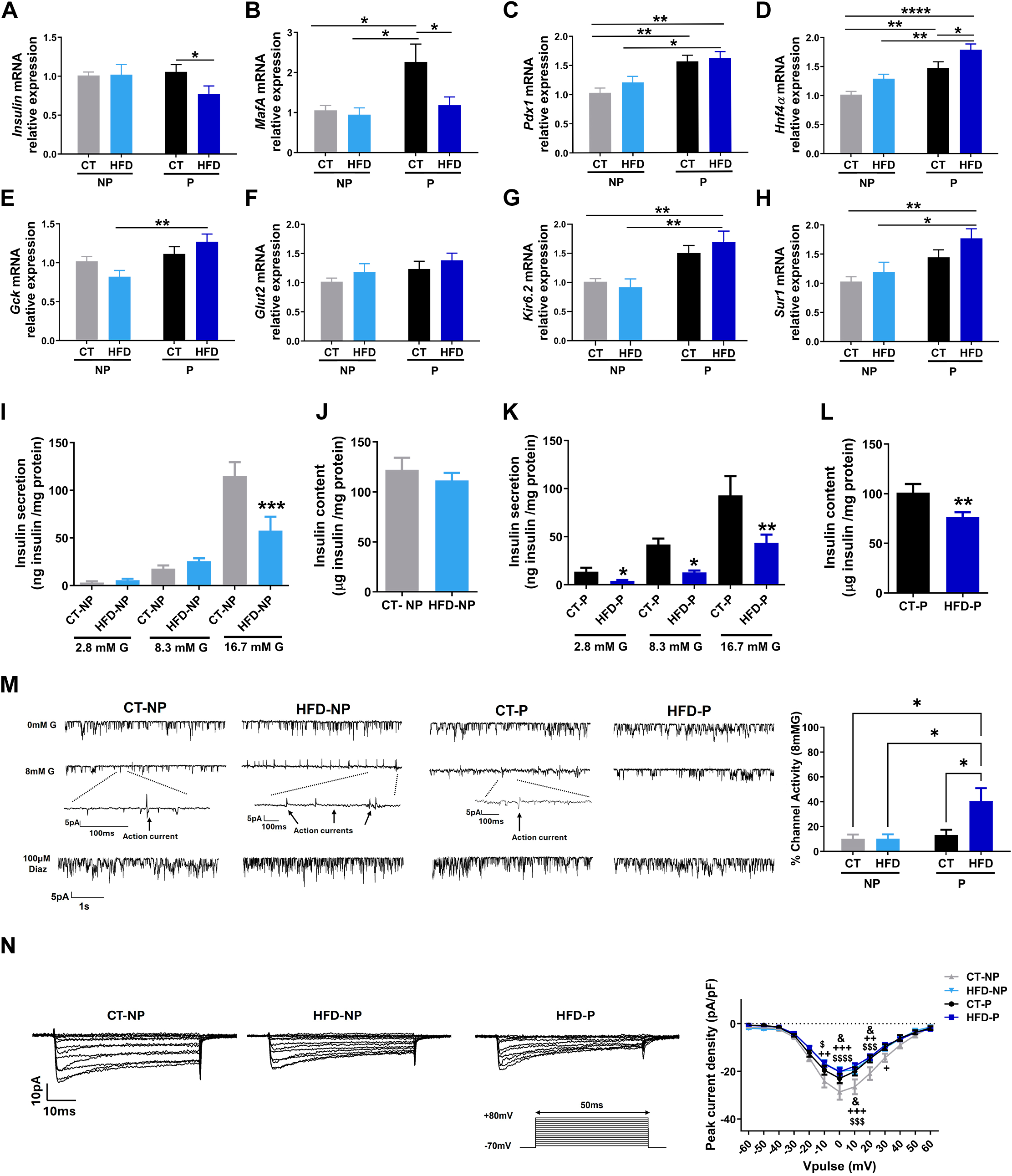
Study of pancreatic β-cell function in control and GDM mice. Islet mRNA levels of **(A)** *Insulin*, **(B)** *Mafa*, **(C)** *Pdx1*, **(D)** *Hnf4α*, **(E)** *Gck*, **(F)** *Glut2*, **(G)** *Kir6.2* and **(H)** *Sur1* from CT-NP (n=10), HFD-NP (n=8), CT-P (n=11) and HFD-P (n= 12-13) mice at GD16 or equivalent length of time. **(I)** Ex vivo glucose-stimulated insulin secretion at 2.8 mM, 8.3 mM and 16.7 mM of glucose in batches of isolated size-matched islets from non-pregnant mice (CT-NP, n= 12-15; HFD-NP, n= 14-15). **(J)** Islet insulin content measurement in batches of islets from non-pregnant mice (CT-NP, n= 42; HFD-NP, n=48). **(K)** Ex vivo glucose-stimulated insulin secretion in batches of isolated size-matched islets from pregnant mice (CT-P, n= 12-15; HFD-P, n=12-13). **(L)** Islet insulin content measurement in batches of islets from pregnant mice (CT-P, n= 43; HFD-P, n=44). Insulin secretion and content were normalized by total islet protein levels. **(M)** Representative recordings of K_ATP_ channel activity in pancreatic β-cells isolated from CT-NP, HFD-NP, CT-P and HFD-P mice. Channel openings are represented by downward deflections, reflecting inward currents due to high K^+^ content of the pipette. The abolition of K_ATP_ channel activity and generations of action currents at 8 mM G was used as a positive control of pancreatic β-cells identity. The graph shows the quantification of the K_ATP_ channel activity in the presence of 8 mM glucose. The effect of glucose was measured after 10 min of acute application in each condition. Data are represented as a percentage of activity with respect to resting conditions (0 mM G). Experiments were carried out at 32-34°C (CT-NP, n= 7; HFD-NP, n=7; CT-P, n= 8; HFD-P, n= 8 cells). **(N)** Voltage-gated Ca^2+^ currents. Representative recordings of voltage-gated Ca^2+^ currents in response to 500 ms depolarizing pulses (−60 mV to +80 mV from a holding potential of −70 mV [inset]). Average relationship between voltage-gated Ca^2+^ current density (currents in pA normalized to cell size in pF) and voltage of pulses (CT-NP, n= 22; HFD-NP, n= 31; CT-P, n= 15; HFD-P, n= 17 cells). Data are presented as means ± SEM. Statistical comparisons were performed using Two-way ANOVA followed by Tukeýs (A-H, M and N) or Sidak’s (I, K) post hoc test. Student’s t test was used in L. *P < 0.05, **P < 0.01 and ****P < 0.001. In (N) $ P < 0.05, $$$ P < 0.001, $$$$ P < 0.0001 (CT-NP vs. HFD-NP); & P < 0.05 (CT-NP vs. CT-P); + P < 0.05, ++ P < 0.01, +++ P < 0.001 (CT-NP vs. HFD-P).

We then analysed pancreatic β-cell function. For this purpose, we measured glucose-stimulated insulin secretion in isolated islets exposed to different glucose concentrations. Islets from HFD-NP mice displayed reduced insulin secretion response to glucose compared to islets from CT-NP although the effect was only significant at the highest glucose concentration (16.7 mM) (Fig. 3I). The decline of insulin secretion capacity was much more evident in islets from HFD-P, which exhibited decreased insulin release in response to both 8.3 and 16.7 mM glucose concentrations compared to CT-P islets (Fig. 3K). In addition, insulin secretion was also found to be downregulated (Fig. 3K). Besides, insulin content was reduced in HFD-P islets (Fig. 3L) while no effect was found in HFD-NP (Fig. 3J).

It is well stablished that the K_ATP_ channel serves as a metabolic sensor playing a central role in insulin secretion. Here, we evaluated K_ATP_ channel activity using the patch clamp technique. We found that pancreatic β-cells from HFD-P mice showed increased activity of the K_ATP_ channel and decreased channel closure in response to glucose compared to CT-P, CT-NP and HFD-NP. This was accompanied with lower occurrence of action currents (Fig. 3M), indicating a reduced action potential firing, which is instrumental in the activation of voltage-gated Ca^2+^ channels and insulin release. No differences were observed between CT-P and CT-NP nor HFD-NP suggesting that the effect was specific for HFD-P β-cells (Fig. 3M). Additionally, voltage-gated Ca^2+^ currents were analysed as they are also a key part of the molecular pathway triggering insulin release. As shown in Fig. 3N, Ca^2+^ currents in response to depolarizing voltages pulses were decreased in HFD-NP b-cells compared to CT-NP cells. Similar effect was observed in HFD-P b-cells (Fig. 3N). In contrast, voltage-gated K^+^ currents were similar in all experimental groups (Fig. S2A-B). The observed alterations in ion channel expression and activity provide an explanation for the diminished insulin secretion observed in mice with GDM.

Next, we examined potential pancreatic β-cell mass changes among the different experimental groups. Morphometric analysis revealed that both pancreatic β-cell area and mass were increased in CT-P and HFD-P vs. CT-NP mice (Fig. 4A-B). Of note, we also found decreased β-cell area in HFD-NP vs. CT-NP (Fig. 4A) but no differences in β-cell mass (Fig. 4B). The number of proliferating pancreatic β-cells, identified by double staining of insulin and Ki67, were increased in both CT-P and HFD-P animals (Fig. 4C). Of note, despite the increment compared to non-pregnant conditions, HFD-P pancreatic β-cells showed a marked decrease in proliferation when compared to CT-P (Fig. 4C).

**Figure 4.**
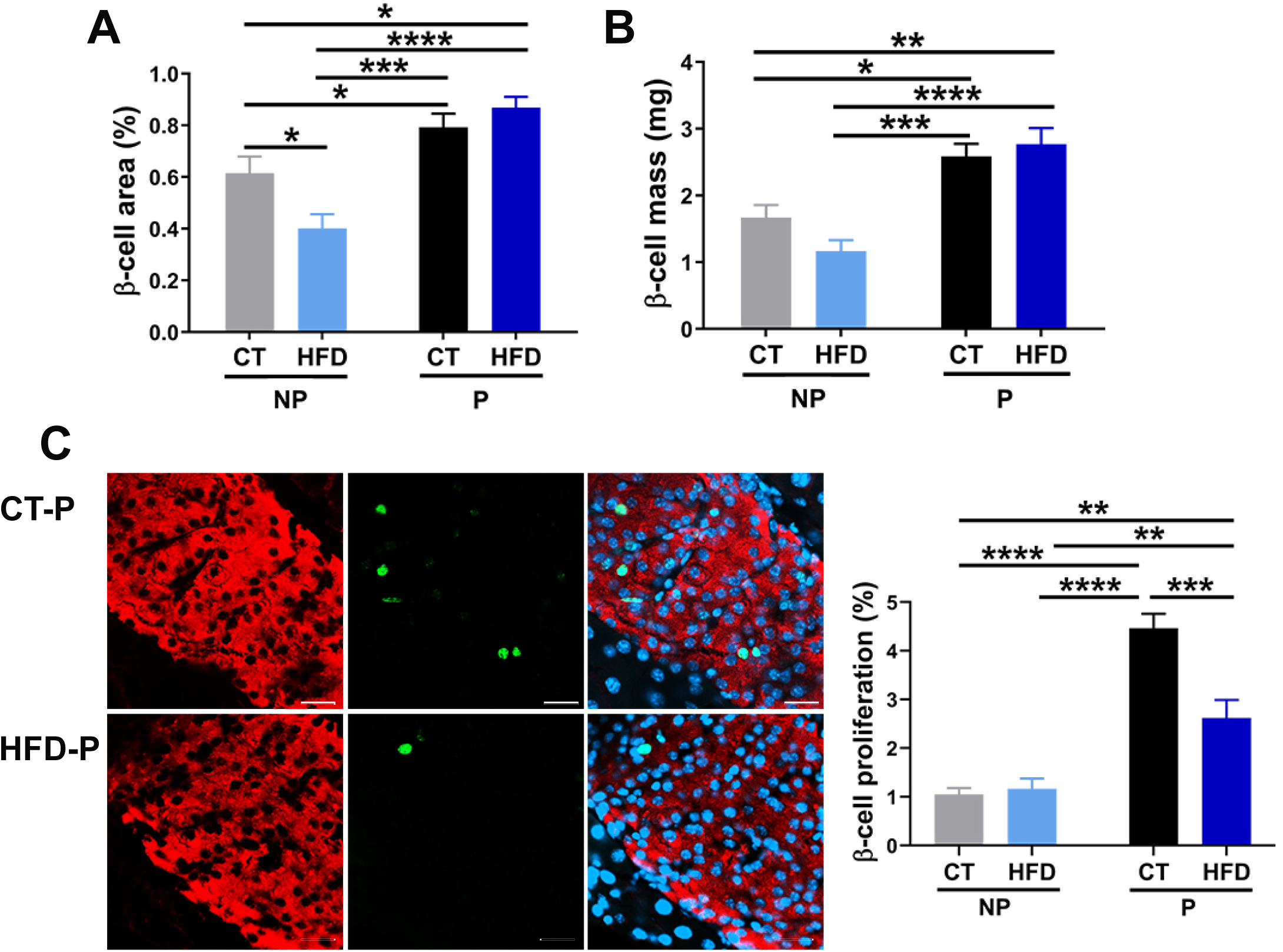
Pancreatic β-cell mass and proliferation in control and GDM mice. **(A)** Quantification of pancreatic β-cell area and **(B)** pancreatic β-cell mass in CT-NP (n = 7), HFD-NP (n=7), CT-P (n= 7) and HFD-P (n=7) at GD15 or equivalent length of time. **(C)** Representative images of pancreas sections stained for insulin (red), Ki67 (green) and nuclei with Hoechst (blue). Scale bar, 20 μm. The graph shows the quantification of pancreatic β-cell proliferation (CT-NP, n= 8; HFD-NP, n= 8; CT-P, n= 7; HFD-P, n= 7). Data are presented as means ± SEM. Statistical comparisons were performed using Two-way ANOVA followed by Tukeýs or Sidak’s post hoc test. *P < 0.05, **P < 0.01, ***P < 0.001 and ****P < 0.0001.

### 3.3. Maternal hormonal changes in GDM

Given the important role of maternal hormones in the adaptive changes and glucose homeostasis control in pregnancy, we evaluated possible alterations in their plasma levels. As it is shown in Fig. 5 we found that placental lactogen and 17 β-estradiol (Fig. 5A-B) were clearly decreased in HFD-P mice compared to their control group. In contrast, progesterone levels were increased in HFD-P (Fig. 5C) while no significant changes in prolactin levels were found (Fig. 5D).

**Figure 5.**
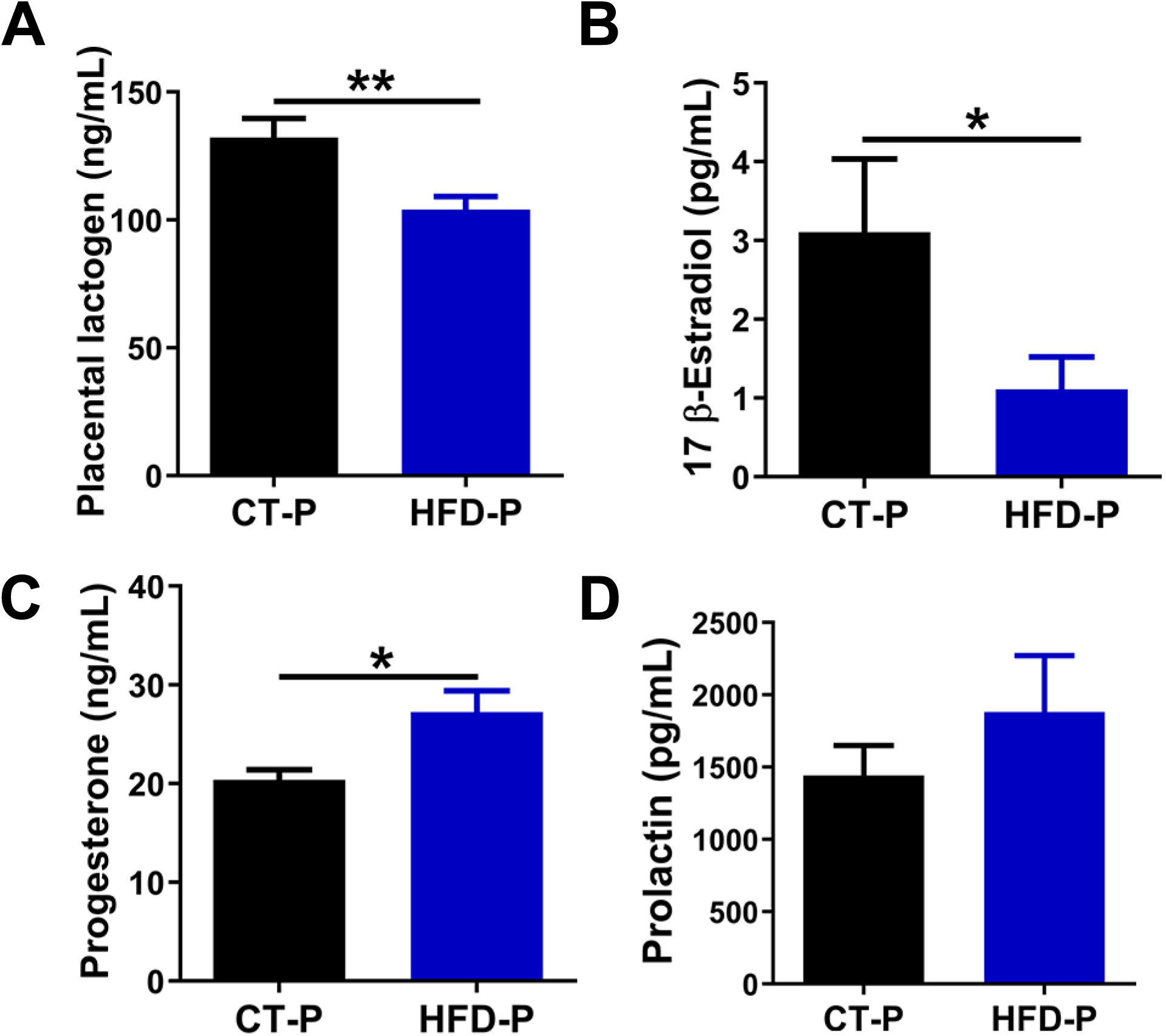
Maternal hormonal changes in GDM mice. Serum levels of **(A)** placental lactogen (CT-P, n=18; HFD-P, n=18), **(B)** 17 β-estradiol (CT-P, n=12; HFD-P, n=12), **(C)** progesterone (CT-P, n=19; HFD-P, n=19) and **(D)** prolactin (CT-P, n= 17; HFD-P, n= 17) measured at GD16. Data are presented as means ± SEM. Statistical comparisons were performed using Student’s t test (A and B) or Mann-Whitney test (C). *P < 0.05, **P < 0.01.

### 3.4. TGFβ signaling in GDM

We quantified the circulating levels of key TGF-β family ligands; TGFβ1, activin-A and inhibin. We found no differences in TGFβ1 serum levels among groups in mice (Fig. 6A). However, a marked increase in activin-A levels was quantified in HFD-P vs. CT-P mice. In contrast, in non-pregnant mice, HFD condition showed the opposite effect with a clear decrease of activin-A in HFD-NP compared to CT-NP (Fig. 6B). A similar situation was found for inhibin levels; a marked increment in the GDM group (HFD-P) compared to its control group (CT-P), and a decline in HFD-NP mice vs. CT-NP. Furthermore, a marked decrease associated with pregnancy was also observed in CT-P compared to CT-NP for inhibin (Fig. 6C). To compare with humans, we measured serum levels of the same TGF-β family ligands in the GDM patients. We observed that TGFβ1 levels were significantly decreased in GDM women (Fig. 6D). In addition to that, activin-A and inhibin concentrations were upregulated in GDM patients (Fig. 6E-F). We also found that BMP-2 levels were enhanced in GDM women despite no statistical significance was reached (Fig. 6G).

**Figure 6.**
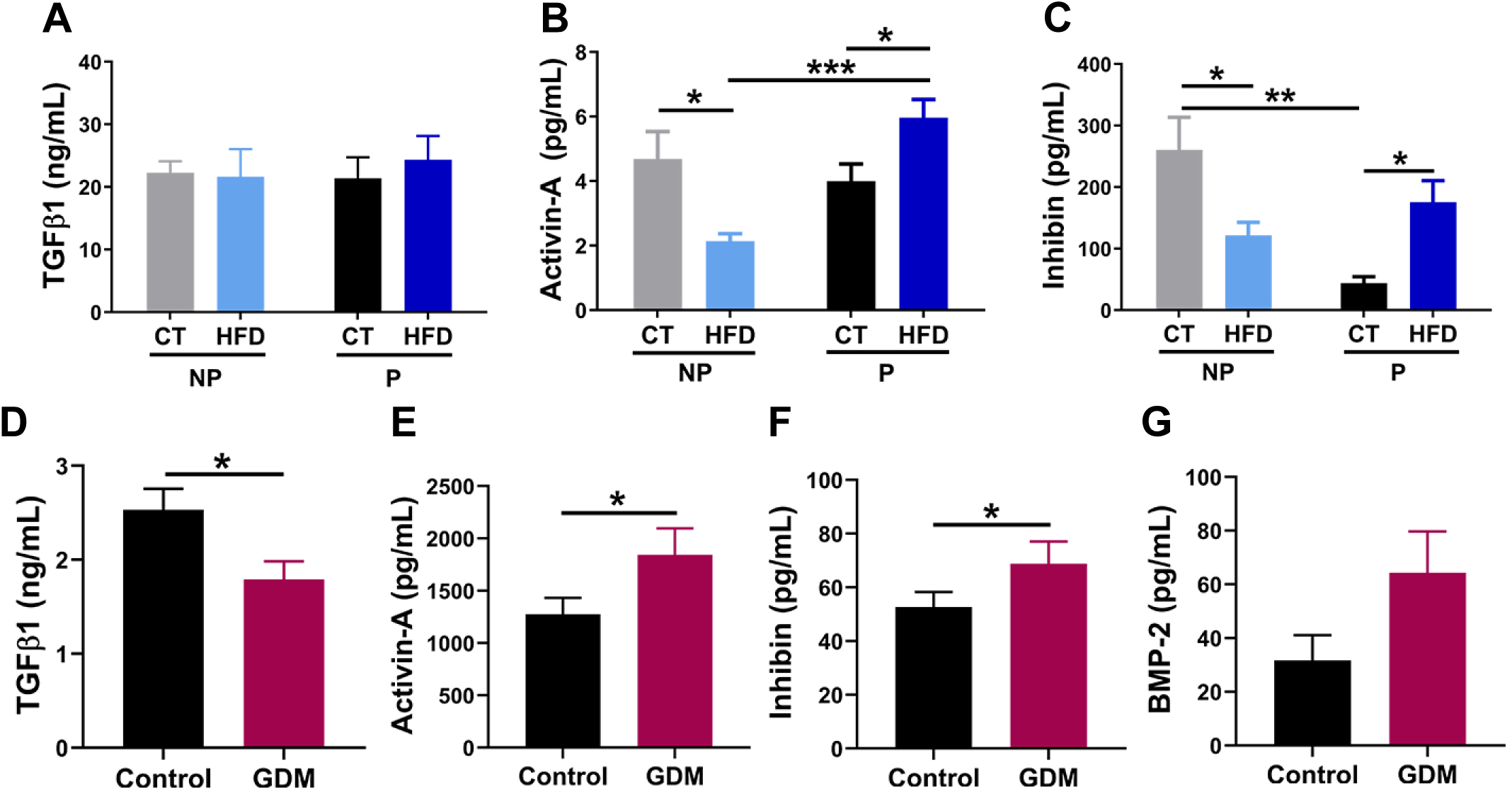
Serum TGFβ ligands in mice and human GDM patients. Mice serum levels of **(A)** TGFβ1 (CT-NP, n=20; HFD-NP, n=13; CT-P, n=21; HFD-P n=19), **(B)** Activin-A (CT-NP, n=22; HFD-NP, n=19; CT-P, n=22; HFD-P n=28) and **(C)** Inhibin (CT-NP, n=18; HFD-NP, n=17; CT-P, n=12; HFD-P n=17). Serum levels of **(D)** TGFβ1, **(E)** Activin-A, **(F)** Inhibin and **(G)** BMP-2 in control and GDM patients (Control (n=10-11); GDM (n= 13-14)). Data are presented as means ± SEM. Statistical comparisons were performed using Two-way ANOVA followed by Sidak’s post hoc test (B, C), Student’s t test (D, E) or Mann Whitney test (F). *P < 0.05, **P < 0.01, ***P < 0.001.

We next evaluated, potential changes in the expression and activity of the main components of the TGFβ/Smad pathway at the endocrine pancreas level in mice. To this end, the islets of Langerhans of the different experimental groups (CT-NP, HFD-NP, CT-P and HFD-P) were isolated, and gene and protein expression assays were conducted.

The most remarkable changes were found in the islets from the GDM group (HFD-P). Increased mRNA expression levels of *TgfβRI* (also known as *Activin receptor-like kinase 5 or Alk5*) and *Smad2* were found in HFD-P islets compared to CT-P (Fig.7A-B). Augmented levels of *Smad3* were also found although the difference was only significant compared to CT-NP (Fig. 7C). Besides, Smad4 was found to be slightly decreased in pregnancy compared to HFD-NP (Fig. 7D). Furthermore, a marked trend of increased *Smad7* expression was found in HFD-P compared to CT-P (Fig. 7E). No significant changes in *Tgfβ1*, *TgfβRII*, *Alk4*, *activin-A (inhba)*, and *inhibin (inha)* gene expression were found in HFD-P vs. CT-P (Fig. S3A-E).

**Figure 7.**
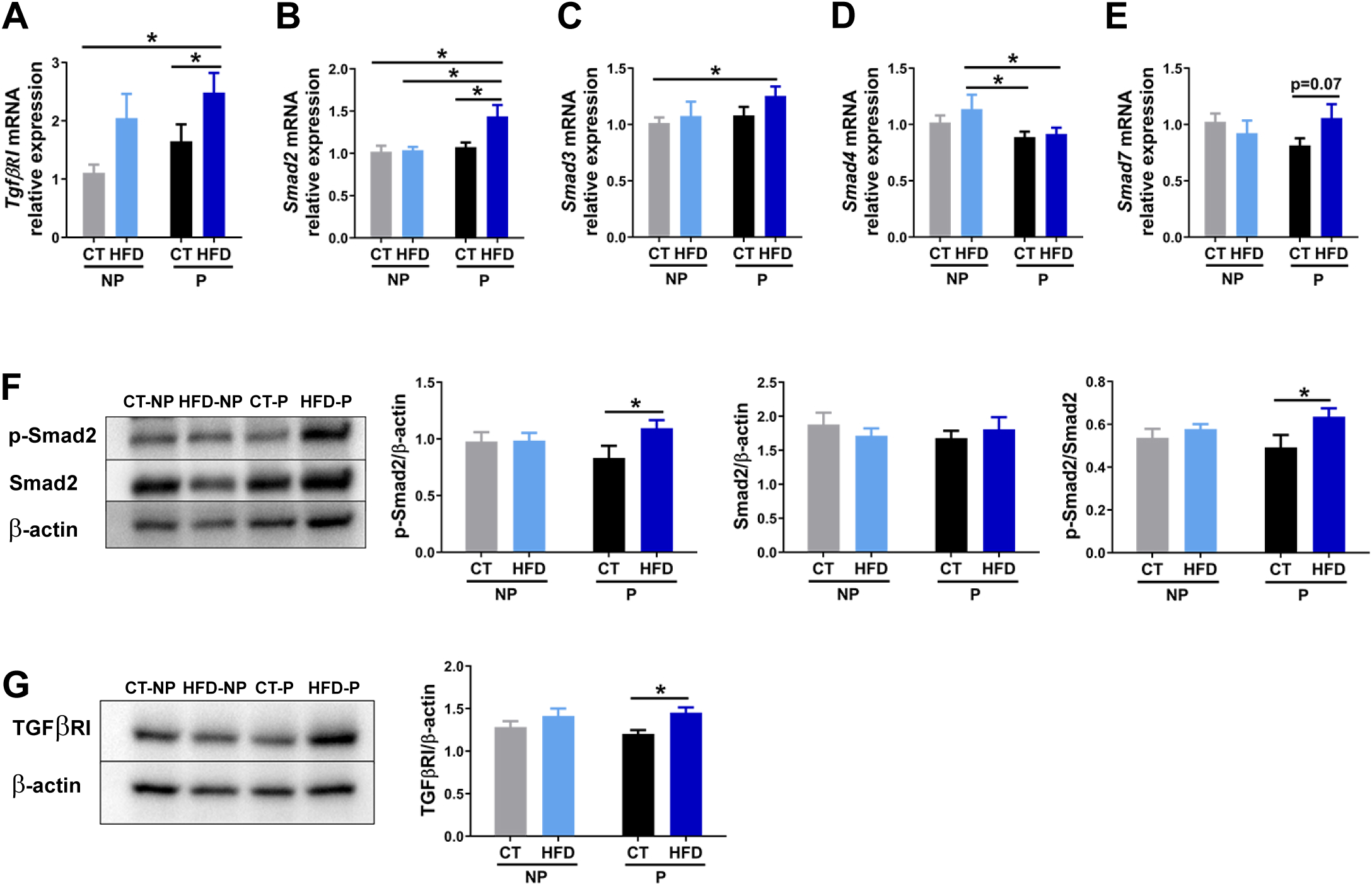
Pancreatic TGFβ/Smad signaling in control and GDM mice. Pancreatic islet mRNA expression levels of **(A)** *TgfβRI*, **(B)** *Smad2*, **(C)** *Smad3*, **(D)** *Smad4* and **(E)** *Smad7* from CT-NP (n=10), HFD-NP (n=8), CT-P (n=11) and HFD-P (n=12-13) mice. Representative western blot images and corresponding quantification showing **(F)** pSmad2, Smad2 and pSmad2/Smad2 protein levels in islets isolated from the different experimental mice groups (CT-NP n= 11; HFD-NP, n= 9; CT-P, n= 8; HFD-P, n= 11), **(G)** TGFβRI (CT-NP, n= 13; HFD-NP, n= 13; CT-P, n= 12; HFD-P, n= 13). All measurements were done at GD16 or equivalent length of time. Data are presented as means ± SEM. Statistical comparisons were performed using Two-way ANOVA followed by Sidak’s (A, B, F and G) or Fisher’s LSD (C, D and E) post hoc test. *P < 0.05.

Then, we examined whether TGFβ/Smad signaling could be differently regulated in GDM (HFD-P group) and/or pregnancy by evaluating changes in the signaling pathway activation at the protein level. Increased phosphorylation of Smad2 as well as elevated pSmad2/Smad2 levels were observed in HDF-P compared to CT-P islets (Fig. 7F). This was in line with an upregulated expression of TGFβRI protein (Fig. 7G). No differences were observed in pSmad3 or pSmad3/Smad3 ratio (Fig. S4A) nor in TGFβRII (Fig. S4B), ALK4 (Fig. S4C) or Smad4 protein expression (Fig. S4D) among groups.

### 3.5. Pharmacological inhibition of TGFβ/Activin-Smad2 signaling in pancreatic islets

As activin-A and inhibin serum levels were found to be elevated in both mice and pregnant women with GDM, it is important to consider the potential implications of these findings. We hypothesized that they may be upstream signaling transducers of pancreatic Smad2 activation. To investigate that, we incubated pancreatic islets in the presence of activin-A and inhibin. We observed that pSmad2 as well as the pSmad2/Smad2 ratio at protein level were dramatically increased compared to controls (Fig. S5A). Then, we assayed the impact of Galunisertib (GL)(LY2157299 monohydrate), a small molecule inhibitor (SMI) of the TGFβRI kinase that specifically downregulates the phosphorylation of Smad2 [24]. We found that GL drastically inhibited the phosphorylation of Smad2 in the presence of the ligands (Fig. 8A). GL also decreased TGFβRI expression under the same experimental conditions (Fig. 8B).

**Figure 8.**
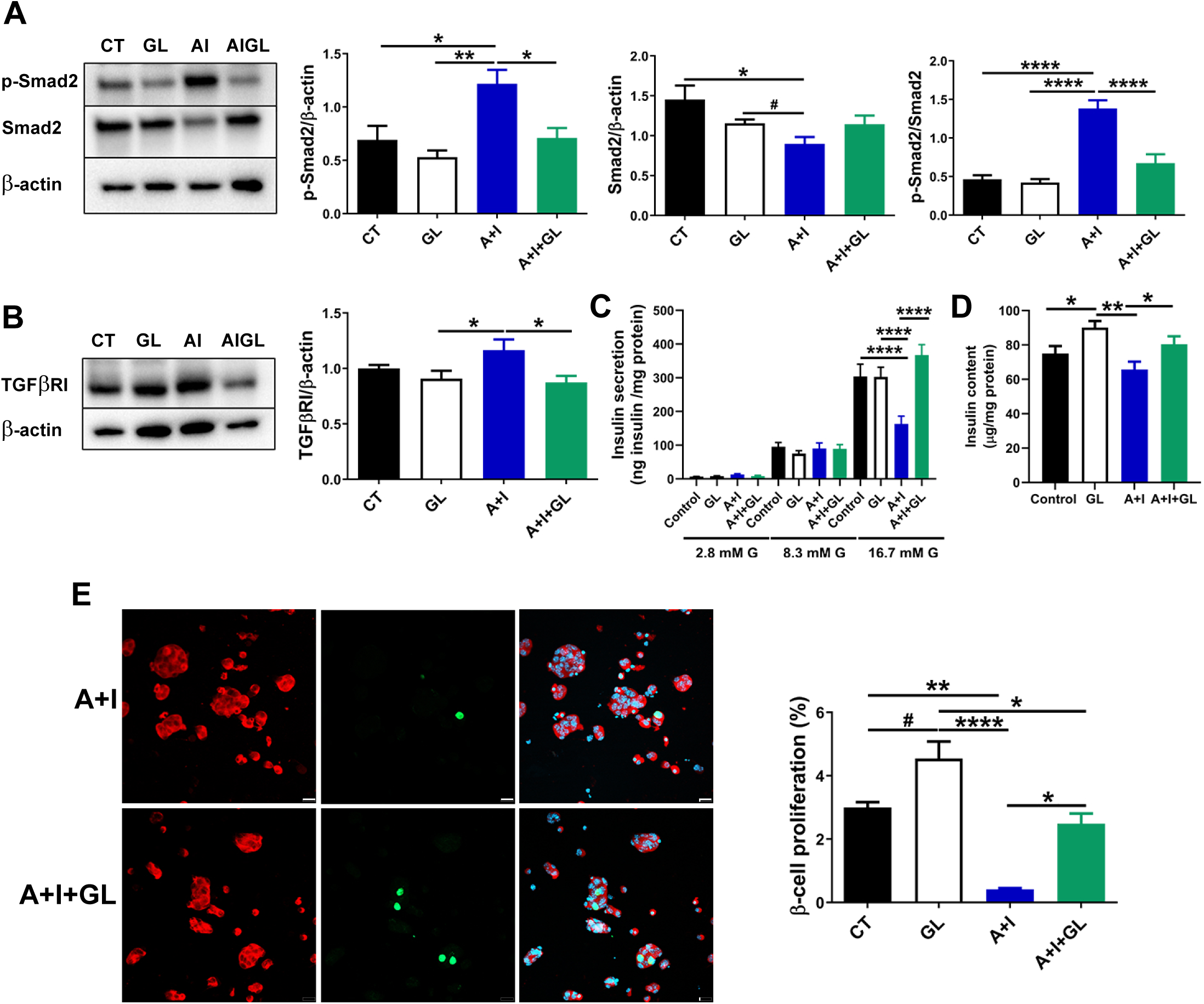
Inhibition of TGFβ/Activin-Smad2 signaling in pancreatic islets of Langerhans. Representative western blot images and corresponding quantification showing **(A)** pSmad2, Smad2 and pSmad2/Smad2 or **(B)** TGFβRI protein levels in islets cultivated for 48 h in the presence of vehicle (CT), galunisertib (GL) (10 μM), activin-A (5 nM) + inhibin (5 nM) (A + I) or activin-A + inhibin + galunisertib (A + I + GL) (n=7-8). **(C)** Ex vivo glucose-stimulated insulin secretion in response to 2.8 mM, 8.3 mM and 16.7 mM of glucose in CT, GL, A+I, and A+I+GL batches of treated islets (n=12-18). **(D)** Insulin content measured in CT, GL, A+I, and A+I+GL batches of treated islets (n=42-49). **(E)** Representative images of islet cells cultured stained for insulin (red), Ki67 (green) and nuclei with Hoechst (blue). Scale bar, 20 μm. The graph shows the quantification of pancreatic β-cell proliferation (CT, n= 6712; GL, n= 6123; A+I, n= 5969 and A+I+GL, n=6559 cells). Data are presented as means ± SEM. Statistical comparisons were performed using One-way ANOVA with Tukeýs post hoc test (A, B and D), Kruskal-Wallis with Dunńs post hoc test (E) or Two-way ANOVA with Tukey’s post hoc test (C). *P < 0.05, **P < 0.01, ****P < 0.0001; #P < 0.05 Student’s t-test (A).

In view of these findings, we wonder whether the TGFβ family ligands could have an impact on pancreatic β-cell function and if the inhibitor GL may act as a modulator of the ligand-induced response. To this end, we quantified glucose-stimulated insulin secretion and found that those islets treated with both activin-A and inhibin showed a marked decrease in insulin release in response to the highest glucose concentration assayed (16.7 mM glucose) (Fig. 8C) and a slight decrease of insulin content (Fig. 8D). Of note, the treatment with GL completely blunted the effect, suggesting a protective effect of GL (Fig. 8C). We also observed that the treatment with both ligands led to diminished pancreatic β-cell proliferation and that the incubation with GL prevented this effect. Of note, the treatment with the inhibitor alone promoted an increase of pancreatic β-cell replication rate (Fig. 8E).

We next explored the individual effects of activin-A and inhibin on TGFβ pathway activation and subsequent Smad2 phosphorylation in pancreatic islets. Islets treated with activin-A for 48 hours showed increased pSmad2 as well as pSmad2/Smad2 ratio. In contrast, inhibin treatment did not have any effect (Fig. S6A). No changes were observed in TGFβRI expression neither in response to activin nor inhibin (Fig. S6B). In addition, we found that GL was able to inhibit the increase in pSmad2 and pSmad2/Smad2 ratio induced by activin-A (Fig. S6C). Decreased expression of TGFβRI was also found when islets were incubated in the presence of activin-A and GL (Fig. S6D). Furthermore, activin-A promoted a marked decrease in pancreatic β-cell insulin secretion (Fig. S6E) and proliferation rate (Fig. S6F), an effect which was abolished by GL (Fig. S6E-F).

## 4. Discussion

The current study first focused on developing a preclinical animal model of GDM that closely resembles the pathophysiology of the disease in women. Our research revealed that HFD treatment three days before and during pregnancy results in a GDM-like phenotype including elevated fasting blood glucose levels, marked glucose intolerance, aggravated insulin resistance, impaired insulin secretion, and compromised β-cell remodelling.

Different animal models have been described in the literature for GDM study. However, they all present a number of limitations to resemble the pathophysiology of the disease. Therefore, GDM modelling remained a challenge when we approached this work. The existing models are based on surgical, chemical, genetic and dietary methods, in an attempt to imitate the GDM features [25–27]. Surgical and chemical methods display significant disadvantages as both evoke severe and irreversible hyperglycaemia associated to acute and extensive pancreatic β-cell death; more similar to T1DM situation [25–27]. Genetic manipulation has also been investigated in understanding GDM disease. However, this approach normally leads to severe impairment of glucose tolerance even in the non-pregnant state, and cannot fully reflect the complex interaction between polygenic and environmental factors, thus limiting their use as specific models for GDM [26].

Nutritional manipulation, including HFD, and high fat and high-sucrose diet (HFHS) diets, has been shown to better mimic the core characteristics of human GDM as well as the metabolic context of the disease, particularly considering that obesity and diet quality are important risk factors for GDM. This is true for rodents but also for larger animal models including sheep and dogs, which may facilitate transfer of findings to clinical management [27]. Furthermore, diet-induced models have also been successful for the study of GDM metabolic consequences for the offspring in the short and long-term [26].

To date, the most common experimental procedures include administration of HFD (45 or 60%) or HFHS for four or six weeks prior mating and during pregnancy. This dietary intervention typically leads to the development of glucose intolerance of varying severity in the mid-to late stages of pregnancy [28–33] and increased fasting insulin levels [30, 32, 34]. However, elevated blood glucose levels and glucose intolerance are observed prior to mating [31, 33], a finding that aligns with our observations following a 4-week treatment with HFD in female mice prior to conception (unpublished data). This is important to consider since perturbed glucose metabolism before the onset of pregnancy does not accurately represent most cases of the human GDM disease but rather reflects a pre-diabetes condition which may be unveiled during pregnancy. By contrast, in our GDM animal model, mice were challenged with HFD (60%) for three days prior mating and throughout pregnancy, with no glucose tolerance impairment ongoing before pregnancy. In line with our results, a study conducted by Pennington and collaborators have shown that acute exposure to HFHS, one week prior to mating and during pregnancy, promoted glucose intolerance and decreased insulin response to glucose at GD17.5 [35]. Of note, when mice were placed on the same diet three weeks before and during pregnancy, the reported GDM alterations were not observed in the mothers leading the authors to highlight the importance of an acute HFHS challenge for GDM development [35].

By using our GDM model we also directly interrogated the status of pancreatic β-cell function and mass in GDM, an aspect that has been less explored in humans due to the difficulty to obtain pancreas samples from women with this disease. We isolated pancreatic islets from GDM mice and found that the capacity of pancreatic β-cells to secrete insulin under all glucose stimulatory conditions assayed was dramatically diminished. Basal insulin secretion was also disrupted as well as insulin content. The decrease in the synthesis and secretion of insulin could be attributable to a combination of different factors. First, we observed that the gene expression of both insulin and Mafa, a key regulator of insulin transcription and other genes important to β-cell function, were markedly decreased in β-cells from GDM mice. The INS gene has been considered as a candidate for the study of GDM in humans. In this regard, the INS-VNTR class III, a polymorphic sequence associated with a less efficient gene transcription of INS gene, has been shown to be more frequent in women who develop GDM [36] although discrepancies can be found depending on the genetic background. Second, we reported that electrical ion channel activity was altered in pancreatic β-cells from GDM mice. In particular, glucose-induced inhibition of K_ATP_ channel activity was impaired, leading to reduced β-cell excitability in response to glucose, which is consistent with the reduced insulin release observed. This finding is of great relevance given the central role of K_ATP_ channel in coupling metabolism and insulin secretion, and that its impaired regulation is a cause of diabetes [37]. Furthermore, voltage-gated Ca^2+^ currents were decreased which also may contribute to the decline of the insulin secretion capacity in the GDM β-cells. These changes, described here for the first time, may help to explain the inability of the β-cells to adapt to the increased insulin demand imposed by pregnancy leading to GDM.

Regarding pancreatic β-cell remodelling, we observed that pancreatic β-cell proliferation rate was clearly increased at GD15 in pregnant compared to non-pregnant mice. It is well described in the literature that proliferation of pancreatic β-cells starts around GD9 with a maximum peak around days GD12-15 [8]. Of note, when we quantified the rate of β-cell proliferation in GDM mice we found that it was markedly lower than that of CT-P mice suggesting that the plasticity of pancreatic β-cells in GDM was reduced. Pancreatic β-cell mass was also augmented in pregnancy compared to non-pregnancy conditions. It is known that during gestation a 3-to 4-fold expansion of β-cell mass occurs, with maximum values around days 15-18 of mouse gestation [10, 38]. Here, we found a 1.6-fold expansion of β-cell mass in CT-P mice at GD15 which leads us to think that the peaking values have not yet been reached but would occur in the following next days. This would be in accordance with the dynamics of pancreatic-β cell mass reported where the proliferation rate peak occurs in advance of the maximal increase in β-cell mass [39].

Maternal hormones play a central role in β-cell adaptation in pregnancy. Here we found decreased placental lactogen and 17 β-estradiol circulating levels while progesterone was increased in GDM mice. This is of relevance since placental lactogen concentration rises during gestation and correlates with enhanced β-cell proliferation and β-cell insulin secretion. The role of estradiol and progesterone in pregnancy is less understood [8] but it is proposed that estradiol increases insulin content and secretion, and exert protective effects against oxidative stress and apoptosis [8, 40]. Likewise, progesterone displays pro-apoptotic effects in β-cells and may contribute to reduce β-cell mass in the peri-partum period [8] and to counterbalance the stimulatory effects of lactogens on β-cells during pregnancy [39].

Once we experimentally demonstrated the suitability of our animal model for GDM study, we interrogated the role of TGFβ/Smad signaling in GDM. To the best of our knowledge, to date, this has not been addressed in the literature. Our studies demonstrate that TGFβRI/pSmad2 signalling was upregulated in GDM pancreatic islets compared to controls. This was evidenced by enhanced Smad2 and TGFβRI mRNA expression levels as well as increased Smad2 activation as indicated by augmented pSmad2 and pSmad2/Smad2 protein levels. Notably, these changes concurred with a decline in pancreatic β-cell function, including decreased insulin secretion and content, altered ion channel activity, and diminished β-cell proliferation. The fact that inhibiting Smad2 signaling abrogated these β-cell alterations suggests that Smad2 could represent an important target for GDM. In agreement with this notion, previous studies have revealed that mice with specific Smad2 ablation in pancreatic β-cells presented a higher β-cell proliferation rate [41, 42], particularly, after pancreatectomy [42]. More recently, it has been demonstrated that pancreatic β-Cell-Specific Smad2 Knockout mice manifested improved glucose tolerance, increased in vivo and ex vivo insulin secretion, and enhanced β-cell mass and proliferation. In addition, they exhibited better glucose tolerance, and augmented insulin secretion and sensitivity when fed a HFD (60%) [17]. The inactivation of Smad2 also promoted differential expression of certain transcription factors which are essential for pancreatic β-cell function and identity [17]. For instance, increased expression of the Mafa gene was identified. In line with these findings, we observed that Smad2 upregulation in GDM was associated with a significant decline in Mafa and insulin gene expression. This could be of relevance at the mechanistic level, as it has been demonstrated that Smad2 is able to physically interact with Mafa leading to inhibit the transcriptional activation of the insulin gene promoter [43]. In a similar manner, TGFβRI knockdown in human pancreatic β-cells upregulated Mafa expression through AKT/FOXO1 pathway and inhibited dedifferentiation [44].

Of note, other studies have shown that Smad3 plays an important function in regulating pancreatic β-cell proliferation, apoptosis and insulin secretion. Deletion of Smad3 has been reported to result in enhanced β-cell proliferation [42, 45] and improved glucose tolerance and insulin secretion [46]. Beneficial effects of Smad3 signaling attenuation in glucose homeostasis has also been observed in db/db [18] and HFD-fed mice [19]. Here we reported, for the first time, elevated pSmad2 signaling in GDM pancreatic islets but no changes in pSmad3, pointing towards a specific role for Smad2 in the impairment of β-cell function and proliferation in GDM.

We demonstrate that circulating levels of activin-A and inhibin were deregulated in GDM, both in our GDM mice model and human GDM patients. Specifically, we observed elevated serum levels of both TGF-β family ligands. This finding is significant for several reasons. First, these molecules may serve as biomarkers reflecting the pathophysiologic mechanisms of GDM involving glucose intolerance and impaired pancreatic β-cell function. In addition, as they are readily obtained from blood samples, they may help to improve the ability to identify women at higher risk of developing GDM and who may benefit from targeted strategies to prevent GDM. Up to date information on TGFβ signaling in GDM is very scarce, with some evidence supporting increased serum maternal levels of activin-A in GDM which were normalized after insulin therapy and restoration of normal blood glucose levels [47]. According to our data, circulating activin-A and inhibin maybe essential upstream signaling transducers of pancreatic Smad2 activation which could be a causal factor of pancreatic β-cell dysfunction and diminished plasticity in GDM.

Previous studies claim that activin-A behaves as a negative regulator of pancreatic β-cell differentiation and function in adult islets. Activin-A treatment significantly decreased mature β-cell genes expression and insulin secretion in MIN6 cells, and mouse and human islets [48]. Others have reported a stimulatory effect of activin-A on insulin secretion in rats [49] and human islets [50, 51]. Contradictory results may be due to the treatment duration, as reduced insulin secretion was reported after a prolonged activin-A treatment of 72 h [48] while acute exposure displayed the opposite effect [49, 50]. In human islets, acute exposure to activin-A also led to decreased insulin secretion, an effect which was reversed by the pre-treatment with the inhibitor follistatin [51]. In agreement with this, our findings support the notion that addition of exogenous activin-A, alone or in combination with inhibin, results in declined insulin secretory capacity of β-cells and decreased proliferation. No previous work has explored the effects of inhibin on pancreatic β-cell function.

Although activin and inhibin in most cases have opposite functions this may depend on the cellular system. For example, both proteins enhance oocyte maturation [52] and blastocyte development [53], with no additive effects [52] or in a synergistic fashion [53]. Our in vitro data show that the combination of activin and inhibin led to Smad2 activation with a direct impact on pancreatic β-cell function and number, and that activin alone displayed a similar response. This may suggest that inhibin neither inhibits nor have an additive effect on that of activin at least under in vitro conditions. Whether or not this occurred in vivo is a complex issue to unravel which merits further investigation as we still have very limited understanding of how the TGFβ family ligands exert their effects on islet pancreatic cells.

Our findings indicate that pancreatic Smad2 activation is upregulated in GDM and that targeting TGFβ/Smad2 signaling may offer novel therapeutic avenues for this pathology. Different approaches have been addressed to block TGFβ signaling including neutralizing antibodies, antisense oligonucleotides and small molecular inhibitors (SMI) [54]. Among SMIs, Galunisertib (LY2157299 monohydrate) is an inhibitor of the TGFβRI kinase which specifically downregulates the phosphorylation of Smad2 and has been moved to clinical investigation given its tolerability and pharmacodynamic profile with encouraging results [24, 55]. Our in vitro studies demonstrate that the treatment with this inhibitor increases pancreatic β-cell proliferation and, even more important, prevent the decrease in the β-cell number found in response to the TGFβ agonists. This could be of clinical relevance if we consider that TGFβRI/Smad2 activation is restraining β-cell mass expansion in GDM. In line with this, growing evidence indicates that SMIs of TGF-β signalling promote increased β-cell replication in both mouse and human β-cells [56, 57] and help to restore mature β-cell identity by inducing expression of β-cell transcription factors [58]. Of note, a recent study has demonstrated that galunisertib is able to rescue apoptosis induced by the loss of GLIS3, a gene which has been associated with type 2, type 1 and neonatal diabetes, in β-like cells [59]. Importantly, our results reflect additional beneficial effects on blocking TGFβRI/Smad2 pathway activation as we found that galunisertib also promoted a marked improvement of pancreatic β-cell function with augmented insulin secretion in response to high glucose.

### 4.1 Conclusions

In the present study, we have developed a suitable preclinical animal model for GDM study which recapitulates the main pathophysiological characteristics of human GDM. Our findings shed light on potential new mechanisms underlying this disease. We found deregulation of serum activin-A and inhibin levels in both mice and human GDM, and highlighted the importance of pancreatic islet Smad2 signaling activation for impaired pancreatic β-cell function and limited regenerative capability of β-cells in GDM. Finally, we propose that attenuation of the TGFβ/Activin-Smad2 signaling could help to alleviate pancreatic β-cell dysfunction and improve plasticity in GDM.

## Funding

This work was supported by grants PID2020-113112RB-I00 (PAM) and PID2023-146795OB-I00 (PAM and IQ) funded by MICIU/AEI/10.13039/501100011033/; ILISABIO22_AP11 Programa de colaboración UMH – Fisabio_ UNISALUT 2022 (PAM and RBB); Generalitat Valenciana CIPROM/2023/27 PROMETEO (PAM and AN).

- CIBER is an initiative of Instituto de Salud Carlos III (Ministerio de Sanidad, Spain).

## Acknowledgment

The authors thank Maria Luisa Navarro, Salomé Ramón, and Beatriz Bonmatí Botella for their excellent technical assistance.

## CRediT authorship contribution statement

**TBB:** Formal analysis, Investigation, Methodology, Validation, Visualization, Writing – review and editing. **HF:** Formal analysis, Investigation, Methodology, Validation, Visualization, Writing – review and editing. **SS:** Formal analysis, Investigation, Methodology, Validation, Visualization, Writing – review and editing. **RBG:** Resources, Funding acquisition, Methodology, Writing – review and editing. **ERG:** Resources, Methodology, Writing – review and editing. **DMB:** Resources, Methodology, Writing – review and editing. **MSS:** Resources, Methodology, Writing – review and editing. **JMP:** Formal analysis, Investigation, Writing – review and editing. **AN:** Resources, Funding acquisition, Writing – review and editing. **IQ:** Formal analysis, Investigation, Methodology, Visualization, Resources, Funding acquisition, Writing – review and editing. **PAM:** Conceptualization, Formal analysis, Funding acquisition, Investigation, Methodology, Project administration, Supervision, Validation, Visualization, Writing – review and editing, Writing – original draft.

## Data availability

The authors confirm that all data are available in the main text or the supplementary materials.

Appendix A. Supplementary material and methods

Appendix B. Supplementary figures (S1-S6)

