## Supplemental material and figures for "Increased TGF β /Activin-Smad2 signaling is associated with pancreatic β -cell dysfunction and glucose intolerance in gestational diabetes mellitus"

**Intraperitoneal Glucose and Insulin Tolerance Tests**

Female mice were fasted for 6 hours (8 am to 2 pm) and then intraperitoneally injected with a glucose load (1.5 g/kg body weight for intraperitoneal glucose tolerance test [IPGTT] and GSIS) or insulin (0.5 UI/kg body weight for non-pregnant and 0.9 UI/kg body weight for pregnant mice for the intraperitoneal insulin tolerance test [IPITT]). Blood was sampled from the tail vein for glucose measurements during IPGTT at 0, 10, 20, 30, 60, 120 minutes and during ITT at 0, 15, 30, 45, 60 minutes. In the IPGTT, blood was collected from the saphenous vein for insulin measurement at 0, 10 and 30 minutes. For glucagon, aprotinin (Sigma-Aldrich, Madrid, Spain) was added to the collection tubes. Glycemia was monitored using an automatic glucometer (Accu-Chek Compact plus; Roche, Madrid, Spain). Plasma insulin and glucagon levels were determined by ELISA (Crystal Chem, Downers Grove, IL, USA). IPGTT and IPITT were performed on gestational day 15 (GD15).

**Mouse serum measurements**

Blood samples for serum analysis were collected at decapitation in non-fasting state at GD16. Hormone concentrations were determined by mouse ELISAs: Insulin (Mercodia, a, Uppsala, Sweden), glucagon, leptin, adiponectin, C-peptide and progesterone (Crystal Chem, Downers Grove, IL, USA), placental lactogen (Biomatik, USA), 17 β-estradiol (Cayman Chemical, USA), and prolactin (Thermo Fisher Scientific).

For serum TGFβ family ligands quantification the following mouse ELISAs were used: TGFβ1, activin-A (Thermo Fisher Scientific), and inhibin (INHA abbexa, UK).

**Islet Isolation and Cell Culture**

Pancreatic islets of Langerhans from females were isolated by collagenase (Sigma, Madrid, Spain) digestion as previously described [1]. The isolation medium contained (in mmol/L): 115 NaCl, 10 $NaHCO_3$, 5 KCl, 1.1 $MgCl_2$, 1.2 $NaH_2PO_4$, 2.5 $CaCl_2$, 25 HEPES, and 5 D-glucose, pH 7.4, as well as 0.25% BSA or, for western blot experiments, 0.1% BSA.

For primary cell culture, isolated islets were dispersed into single cells by trypsin enzymatic digestion. Cells were centrifuged and resuspended in RPMI 1640 without phenol-red (Gibco, Thermo Fisher Scientific Inc., Carlsbad, CA) and with 10% charcoal dextran treated serum (Hyclone, USA), 2 mM glutamine, 100 U/mL penicillin and 0.1 mg/mL streptomycin. Cells were then planted on glass covers and cultured at 37ºC in a humidified atmosphere of 95% $O_2$ and 5% $CO_2$ for 24 hours.

For in vitro experiments, isolated islets or single cells were cultured at 37ºC in a humidified atmosphere of 95% $O_2$ and 5% $CO_2$ for 48 hours and treated with vehicle, 5 nM Activin A (Biotechne, UK) and/or 5 nM Inhibin-A (Biotechne, UK) and/or 10 µM Galunisertib (GL) (MedChemExpress, USA).

**Ex vivo insulin secretion and content measurements**

Freshly isolated islets were left to recover in the isolation medium at 37ºC for 2 hours. Afterward, batches of 5 size-matched islets were transferred to a 24-well plate with 400 µL of a solution containing (secretion buffer) (in mM) 140 NaCl, 4.5 KCl, 2.5 $CaCl_2$, 1 $MgCl_2$, 20 HEPES, and 2.8 mmol/L glucose for 1 hour (2 hours when islets were previously cultured for 48 hours), pH = 7.35, and left at 37ºC and 5% $CO_2$. Then islets were transferred to 400 µL of secretion buffer containing either 2.8, 8.3 or 16.7 mmol/L glucose for 1 hour. Afterwards, supernatants were collected and frozen for subsequent insulin secretion measurements. For insulin content measurements, the islets were handpicked and transferred to 20 µL of ethanol/HCl buffer and incubated overnight at 4°C. The supernatant fraction was collected for insulin content measurement. Insulin secretion and content were normalized by protein content and measured using mouse ELISA (Mercodia). Protein concentration was measured by the Bradford dye method.

**Patch Clamp recordings**

$K_{ATP}$ channel activity was recorded using standard patch-clamp recording procedures in isolated pancreatic β-cells as previously described [2]. Around 80–90% of the single cells were identified as β-cells by their response to high glucose, consisting in action currents in the cell-attached configuration. For the patch-clamp recordings of voltage-gated $K^+$ and $Ca^{2+}$ currents the standard whole-cell configuration was used. For $K_{ATP}$ channel activity and voltage-gated $K^+$ currents, bath solution contained (in mM): 5 KCl, 135 NaCl, 2.5 $CaCl_2$, 10 HEPES and 1.1 $MgCl_2$ and was supplemented with glucose as indicated (pH 7.4 with NaOH). For voltage-gated $Ca^{2+}$ currents bath solution contained (mM): 118mM NaCl, 20mM TEA-Cl, 5.6mM $CaCl_2$, 1.2mM $MgCl_2$, 5mM HEPES and 5mM glucose (pH: 7.4 with NaOH). For recording $K_{ATP}$ channel activity the pipette solution contained (in mM): 140 KCl, 1 $MgCl_2$, 10 HEPES and 1 EGTA (pH 7.2). For recordings of voltage-gated $K^+$ currents, the pipette was filled with the following: 120 mM KCl, 1 mM $MgCl_2$, 1 mM $CaCl_2$, 3 mM MgATP, 10 mM EGTA, and 10 mM HEPES (pH: 7.15 with KOH). A similar medium was used for the $Ca^{2+}$ current measurements except that KCl was equimolarly replaced by CsCl and pH adjusted with CsOH (pH: 7.20 with CsOH).

$K^+$ currents were recorded in response to depolarizing voltage pulses of −60 mV to +80 mV from a holding potential of −70 mV. $Ca^{2+}$ currents were recorded in response to depolarizing voltage pulses of −60 mV to +60 mV from a holding potential of −70 mV. For $K^+$ and $Ca^{2+}$ currents density quantification, $K^+$ and $Ca^{2+}$ currents (in pA) were normalized to the cell capacitance (in pF).

**RNA Isolation and Gene Expression Analysis by Real-Time Quantitative PCR**

Total islet RNA was extracted with RNeasy Micro Kit (Qiagen, Madrid, Spain) according to the manufacturer's instructions and quantified with a Nanodrop 2000 (Thermo Fisher Scientific, Waltham, MA), followed by cDNA synthesis with high-capacity cDNA reverse transcription kit (Applied Biosystems, Foster City, CA). Quantitative real-time polymerase chain reaction (qRT-PCR) assays were carried out in a final volume of 10 µL, containing 200 nM of each primer, 1 µL of cDNA and 1 × IQ SYBR Green Supermix (Bio-Rad, Hercules, CA), in a CFX96 Real Time System (Bio-Rad). The resulting values were analysed with CFX Maestro 2.3 (Bio-Rad) and expressed as the relative expression respect to control levels using the comparative 2-ΔΔCT method. HPRT ((Hypoxanthine-guanine phosphoribosyl transferase) was used as the housekeeping gene. Further information on primer sequences is described in Supplemental Table 1.

**Western blotting**

Islets were washed with cold PBS and lysed in Laemmli buffer. Proteins were separated on precast gels (8-16% Mini-Protean TGX Bio-Rad) and subsequently transferred to polyvinylidene disulfide membranes PVDF (Amersham-Cytiva). Then, membranes were blocked in Tris-buffered saline/Tween (TBST) buffer containing 5% low-fat milk protein for 1 hour at room temperature. Blots were then incubated overnight at 4ºC with the following primary antibodies: anti-pSmad2 (ab188334, Abcam, 1:500), anti-Smad2 (D43B4, Cell Signaling, 1:1000), anti-TGFβRI (ab235578, Abcam, 1:500) and anti-βActin (A5316, Sigma, 1:5000). Also, anti-p-Smad3 (sc-517575, Santa Cruz, 1:500), anti-smad3 (sc-101154, Santa Cruz, 1:200), anti-TGFβRII (ab61213, Abcam, 1:500), anti-Smad4 (ab40759, Abcam, 1:2000) or ALK4 (ARG40270, Arigo Biolaboratories, 1:1000) as shown in supplemental figures. Afterward, the membranes were washed with TBST buffer and incubated for 1 hour at room temperature with appropriate secondary peroxidase-conjugated antibodies (Bio-Rad, Richmond, CA, USA; 1:5000). Membranes were developed using SuperSignal West Femto chemiluminescent reagent (Thermo Scientific, Rockford, IL, USA), and visualized with the BioRad ChemiDoc XRS+ System (Bio-Rad Laboratories). The intensity of the bands was quantified using Image Lab software (version 4.1, Bio-Rad Laboratories).

**β-Cell Mass and Proliferation Analysis in pancreatic sections**

Pancreas were removed at GD15 or equivalent length of time for non-pregnant mice, weighed and fixed in ice-cold 4% paraformaldehyde overnight at 4ºC. Then pancreata were embedded in paraffin, and tissue sections (5 µm) were prepared. Antigen retrieval was performed by heating the samples for 20 minutes in citrate buffer (10 mM, pH 6.0), and blocking was done by incubation in phosphate buffered saline containing 1% bovine serum albumin, 0.1% triton-X100, 10% goat serum at room temperature for 2 h. After washing, samples were incubated overnight at 4°C with anti-insulin antibody (8138, Cell Signaling, 1:400), washed, and, then incubated with goat anti-mouse Alexa Fluor 594 secondary antibody (A32742, Life Technologies, Carlsbad, CA, 1:500) for 1 hour at room temperature. Nuclei were stained with 10 µg/mL Hoechst (Invitrogen, Barcelona, Spain). Samples were mounted with Fluoromont-G (Life Technologies, Carlsbad, CA). Images were acquired with an IN Cell Analyzer 6000 system (GE Healthcare, Little Chalfont, UK). Each slide was completely scanned by capturing images from no overlapping fields. The total pancreatic area and insulin area was quantified with the ImageJ software (National Institutes of Health, Bethesda, MD). Quantification was done on at least three sections per pancreas, separated by 200 µm, from each animal. β-cell area was calculated by measuring the total insulin-stained area normalized by the total pancreatic area. β-cell mass was calculated by multiplying the β-cell area by the pancreas weight [3].

For the analysis of β-Cell proliferation, pancreas sections were incubated overnight at 4°C in the presence of the primary antibodies (anti-insulin antibody (8138, Cell Signalling, Danvers, MA, USA, 1:400) and anti-Ki67 antibody (12202, Cell Signalling Technology, Danvers, MS, 1:225)) and, subsequently, with goat anti-mouse Alexa Fluor 594 (A32742, Life Technologies) and goat anti-rabbit 488 (A32731, Life Technologies) secondary antibodies (1:500) dilution for 1 hour at room temperature. Nuclei were stained with 10 µg/mL Hoechst (Invitrogen). Slides were mounted with Fluoromont-G (Life Technologies, Carlsbad, CA). Images were acquired with a LSM900 confocal microscope at 40x. Ki-67-positive nuclei were scored only in cells that were also positive for insulin. Quantification analysis was performed using the ImageJ software (National Institutes of Health, Bethesda, MD).

**β-Cell Proliferation analysis in primary islet cell culture**

Cells were fixed with 4% paraformaldehyde for 10 minutes at room temperature and washed with PBS. Permeabilization was done with 0.5% Triton X-100 for 3 min. Non-specific interactions were blocked with 1% bovine serum albumin, 0.1% Triton-X100, 10% goat serum in phosphate buffered saline for 1 hour at room temperature. Cells were then incubated with anti-insulin antibody (8138, Cell Signalling, Danvers, MA, USA, 1:400) and anti-Ki67 antibody (12202, Cell Signalling Technology, Danvers, MS, 1:100) overnight at 4ºC. After washing, cells were incubated with the secondary antibodies goat anti-mouse Alexa Fluor 594 and goat anti-rabbit 488 both for 1 hour at room temperature. The nuclei were stained with 10 µg/mL Hoechst (Invitrogen, Barcelona, Spain) for 10 min at room temperature. Samples were mounted using Fluoromont-G (Life Technologies, Carlsbad, CA). Images were acquired with a Zeiss LSM900 confocal microscope at 20x. Proliferation rate was expressed as the percentage of β-cells with Ki67-positive nuclei respect to the total number of β-cells. Quantification analysis was performed with the ImageJ software (National Institutes of Health, Bethesda, MD).

**Clinical patient data, sample collection and measurements**

Patients were recruited at the Hospital Vinalopó (Elche, Spain). Women were screened for GDM at 24-28 weeks following the Spanish Diabetes and Pregnancy Group recommendations. Subjects were challenged with a 1-hour 50-g glucose challenge test, if values were ≥ 140 mg/dL, they underwent a 3-hour 100-g oral glucose tolerance test (OGTT). GDM was diagnosed if two or more plasma glucose levels met or exceeded the threshold according to National Diabetes Data Group and 3rd Workshop-Conference on Gestational Diabetes [4]. Women were excluded if they had pre-existing diabetes, prior gestational diabetes or multifetal gestation; or on medication which is known to affect carbohydrate metabolism during pregnancy. The Ethics Committee of the Hospital approved the experimental protocol (identification code: BIODIAGES; acceptance date: 26 january 2023). All participants gave their written, informed consent.

Blood samples were obtained at fasting between 27 and 29 weeks of pregnancy for biochemistry analysis. Plasma insulin and C-peptide were determined by chemiluminescence on an Atellica IM analyzer (Siemens Healthineers). Triglycerides were determined by spectrophotometric method (Atellica IM analyser, Siemens Healthineers). Human ELISAs kits were used to quantify Adiponectin (Crystal Chem, Downers Grove, IL, USA), TNF$\alpha$, IL-6, TGF$\beta$1 and BMP-2 (Invitrogen), activin-A (R&D Systems, Inc, USA) and inhibin (Raybiotech, USA).

| GEN | FORWARD (5'->3') | REVERSE (5'->3') |
|---|---|---|
| *Hprt* | GGTTAAGCAGTACAGCCCCA | TCCAACACTTCGAGAGGTCC |
| *Ins* | TTATTGTTTCAACATGGCCC | CAAAGGTGCTGCTTGACAAA |
| *Pdx1* | GGCCTGGAAGAGCCCAACCG | TGTGTAAGCACCTCCTGCCCACT |
| *Mafa* | CATCCGACTGAAACAGAAG | ATTTCTCCTTGTACAGGTCC |
| *Hnf4α* | TCTGGATGACCAGGTGGCGCT | GGACACACGGCTCATCTCCGC |
| *Gck* | TTCAGCTTCTGGCCTCCCACAG | AAAACAGCCAGGTCTGGGCAGC |
| *Glut2* | ATCGCTCCAACCACACTCAG | CTGAGGCCAGCAATCTGACTA |
| *Kir6.2* | CCCGTCTTGGCCCTCCATGATT | TATCCCGCCGTGTCTGTGTGG |
| *Sur1* | TGCCTCAGGACAAGCAACC | GACCACTGTCCTCTTGTCATC |
| *TgfβRI (activin receptor-like kinase 5 or Alk5)* | TGGATCAGGTTTACCACTGCT | CAACTTCTTCTCCCCACCAT |
| *TgfβRII* | TAACAGTGATGTCATGGCCAGCG | AGACTTCATGCGGCTTCTCACAGA |
| *Alk4 (activin A receptor type 1B)* | CAGAGTTATGAGGCCTTGCG | AGAGTCTTCTTGATGCGCAGA |
| *Smad2* | AAGCCATCACCACTCAGAATTG | CACTGATCTACCGTATTTGCTGT |
| *Smad3* | AGGGGCTCCCTCACGTTATC | CATGGCCCGTAATTCATGGTG |
| *Smad4* | GCAGCTCTTGGATGAAGTCC | GGCAGCAAACACATCTCTCA |
| *Smad7* | GCAGGCTGTCCAGATGCTGT | GATCCCCAGGCTCCAGAAGA |
| *Tgfβ1* | TGCCCCTATATTTGGAGCCTG | GTAGTAGACGATGGGCAGTGG |
| *Activin-A (Inhba)* | TGGTGCCAGTCTAGTGCTTC | CCGTCACTCCCATCTTTCTT |
| *Inhibin (Inha)* | TCCTTTTGCTGTTGACCCTA | CCCCAAGGCATCTAGGAATA |

Supplemental Table 1. List of primers.

#### Supplementary Figures

### Increased TGF β /Activin-Smad2 signaling is associated with pancreatic β -cell dysfunction and glucose intolerance in gestational diabetes mellitus

Talía Boronat-Belda[1]*, Hilda Ferrero[1,2]*, Sergi Soriano[1,2], Elena Ribes-García[3], Rubén Betoret-Gustems[3], Daniel Martínez-Bañón[4], Mónica Serrano-Selva[3], Juan Martínez-Pinna[1,2], Angel Nadal[1,5], Iván Quesada[1,5] and Paloma Alonso-Magdalena[1,5]

[1] Instituto de Investigación, Desarrollo e Innovación en Biotecnología Sanitaria de Elche (IDiBE), Universidad Miguel Hernández de Elche, Elche, Spain.
[2] Departamento de Fisiología, Genética y Microbiología, Universidad de Alicante, Alicante, Spain.
[3] Servicio de Ginecología y Obstetricia, Hospital Universitario del Vinalopó, Elche, Spain
[4] Laboratorio Ribera Lab Hospital Universitario del Vinalopó, Elche, Spain
[5] CIBER de Diabetes y Enfermedades Metabólicas Asociadas (CIBERDEM), Instituto de Salud Carlos III.

*These authors contributed equally to this work

**Corresponding author**

Paloma Alonso-Magdalena, Instituto de Investigación, Desarrollo e Innovación en Biotecnología Sanitaria de Elche (IDiBE), Universidad Miguel Hernández de Elche, 03202-Elche, Alicante, Spain.

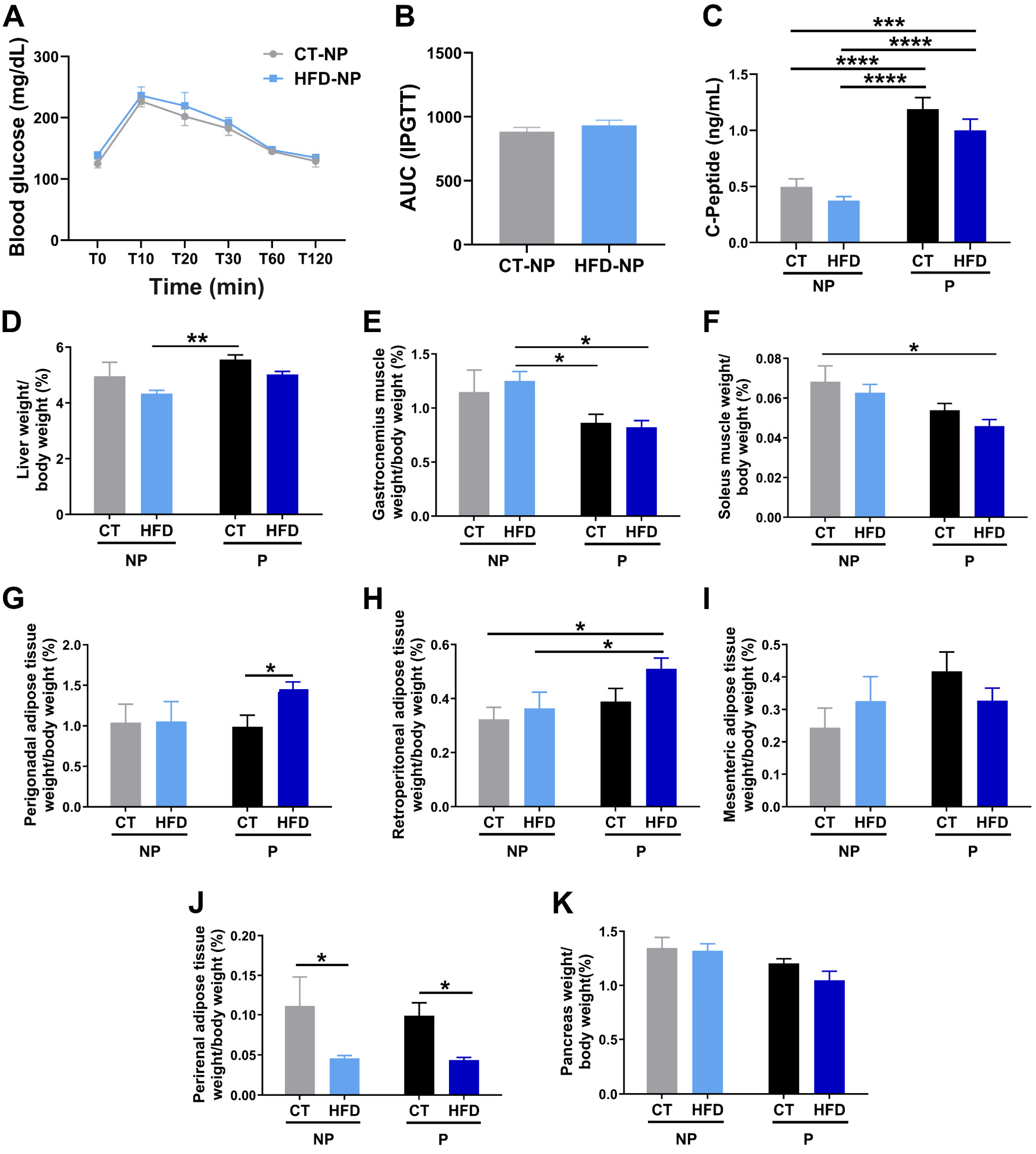


**Supplemental Figure 1. (A)** Intraperitoneal glucose tolerance test (IPGTT) in mice fed with control diet (CT-NP, n=6) or HFD (HFD-NP, n=5) during 3 days. **(B)** Area under the curve (AUC) from the IPGTT. **(C)** C-peptide serum levels in fed state measured at GD16 or equivalent length of time (CT-NP, n=16; HFD-NP, n=19; CT-P, n=19; HFD-P, n=19). **(D)** Liver, **(E)** gastrocnemius muscle, **(F)** soleus muscle, **(G)** perigonadal adipose tissue, **(H)** retroperitoneal adipose, **(I)** mesenteric adipose, **(J)** perirenal adipose and **(K)** pancreas weight tissue normalized by body weight from CT-NP, n=4-10; HFD-NP, n=5-10; CT-P, n=7-8; HFD-P, n=6-8. Data are presented as means ± SEM. Statistical comparisons were performed using Two-way ANOVA followed by Tukey´s (C, D, E and F) or Fisher's LSD (G, H and J) post hoc test. *P < 0.05, **P < 0.01, ***P < 0.001, ****P < 0.0001.

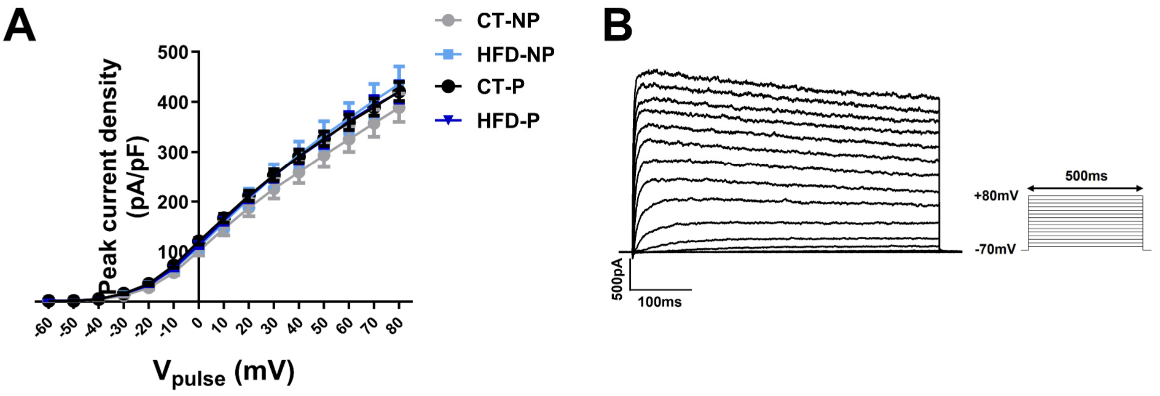


**Supplemental Figure 2. (A)** Voltage-gated $K^+$ currents. Average relationship between voltage-gated $K^+$ current density (currents in pA normalized to cell size in pF) and voltage of pulses. **(B)** Representative recordings of voltage-gated $K^+$ currents in response to 500 ms depolarizing pulses (−60 mV to +80 mV from a holding potential of −70 mV [inset]) in CT-NP, n= 16; HFD-NP, n= 20; CT-P, n= 37; HFD-P, n= 47 cells). Data are presented as means ± SEM.

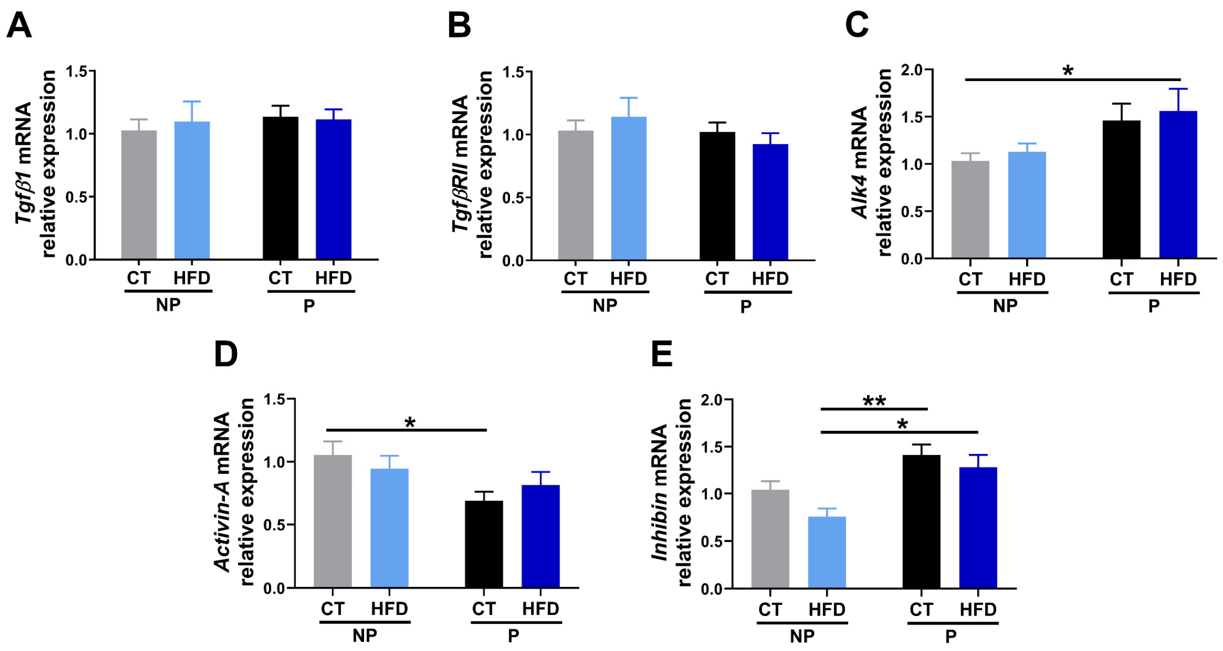


**Supplemental Figure 3.** Islet mRNA levels of **(A)** *Tgfβ1*, **(B)** *TgfβRII*, **(C)** *Alk4*, **(D)** *Activin-A*, **(E)** *Inhibin* from CT-NP, n=10; HFD-NP, n=8; CT-P, n=11; HFD-P, n=12-13 mice at GD16 or equivalent length of time. Data are presented as means ± SEM. Statistical comparisons were performed using Two-way ANOVA followed by Tukey´s (E) or Fisher's LSD (C and D) post hoc test. *P < 0.05, **P < 0.01.

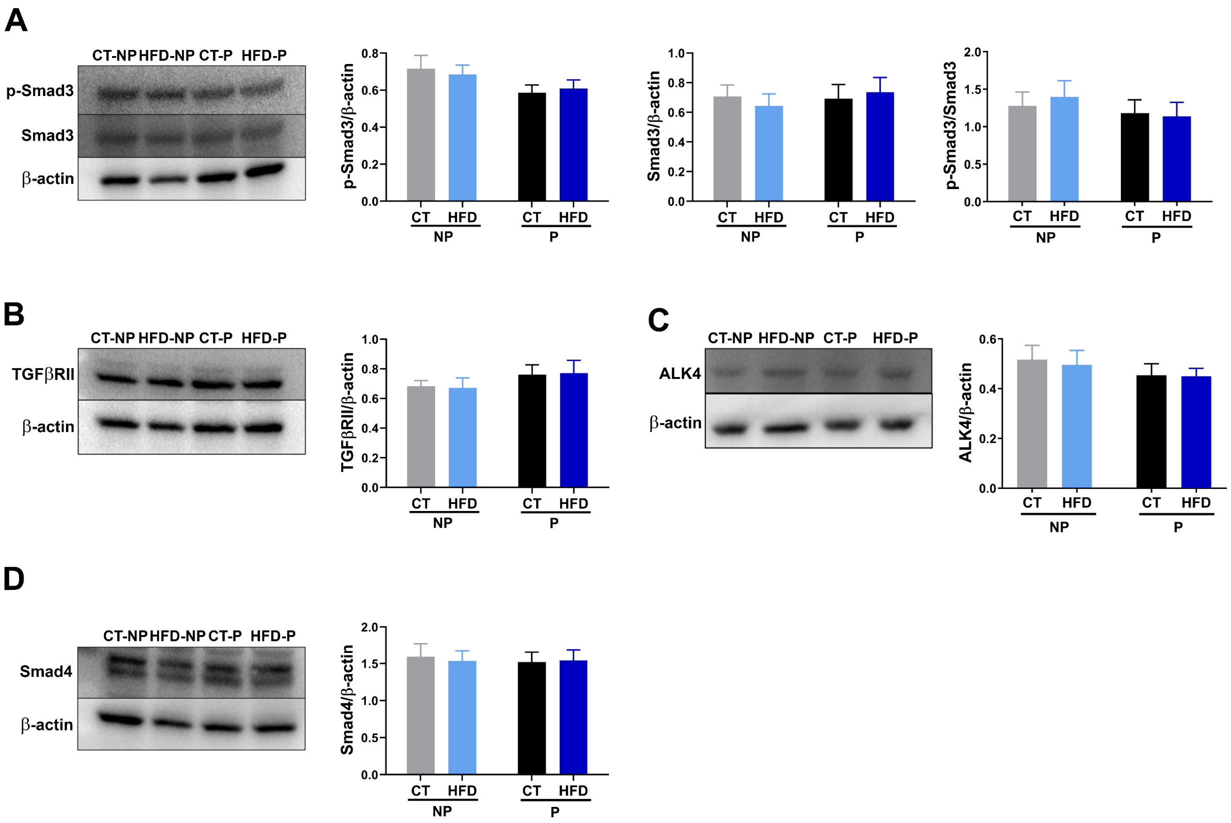


**Supplemental Figure 4.** Representative western blot images and corresponding quantification showing **(A)** pSmad3, Smad3 and pSmad3/Smad3 (CT-NP, n= 18; HFD-NP, n= 16; CT-P, n= 17; HFD-P, n= 18), **(B)** TGFβRII (CT-NP, n= 12; HFD-NP, n= 12; CT-P, n= 12; HFD-P, n= 12), **(C)** ALK4 (CT-NP, n= 12; HFD-NP, n= 12; CT-P, n= 12; HFD-P, n= 12) and **(D)** Smad4 (CT-NP, n= 10; HFD-NP, n= 10; CT-P, n= 11; HFD-P, n= 11) protein levels in islets isolated from the different experimental groups. Data are presented as means ± SEM.

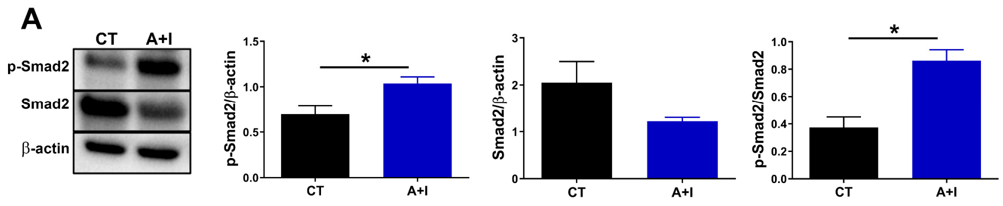


**Supplemental Figure 5. (A)** Representative western blot images and corresponding quantification showing pSmad2, Smad2 and pSmad2/Smad2 protein levels (n=4). Data are presented as means ± SEM. Statistical comparisons were performed using Mann-Whitney test. *P < 0.05

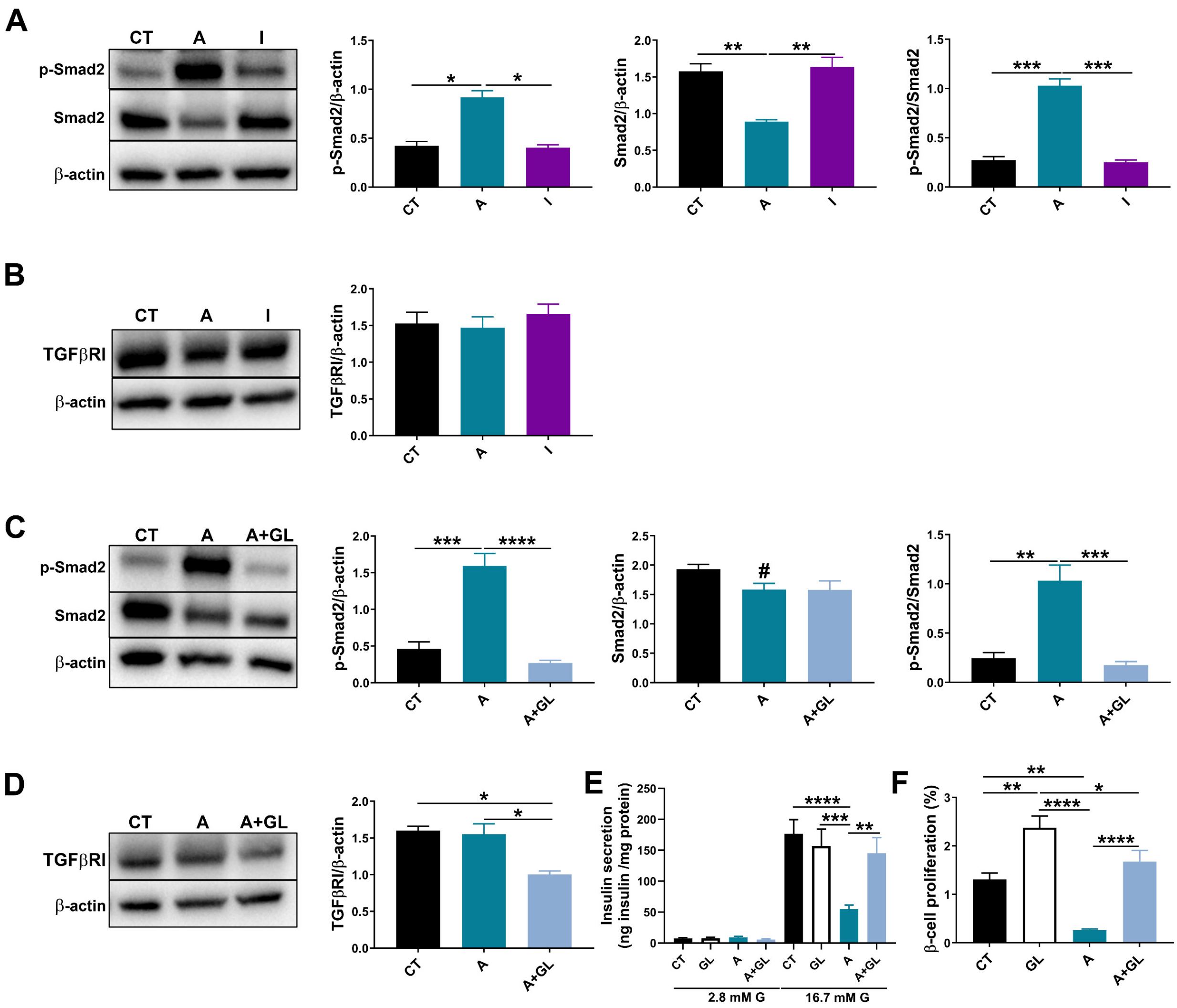


**Supplemental Figure 6.** Representative western blot images and corresponding quantification showing **(A)** pSmad2, Smad2 and pSmad2/Smad2, **(B)** TGFβRI protein levels in islets treated for 48 h with CT, A or I (n=4). Representative western blot images and corresponding quantification showing **(C)** pSmad2, Smad2 and pSmad2/Smad2 and **(D)** TGFβRI in islets treated for 48 h with CT, A or A+GL (n=4). **(E)** Glucose stimulated insulin secretion in batched of islets treated for 48 h with CT, GL, A or A+GL (n=8-12) **(F)** Pancreatic β-cell proliferation (CT, n=5995; GL, n= 6013; A, n= 6415; and A+GL, n=6036 cells). Data are presented as means ± SEM. Statistical comparisons were performed using One-way ANOVA followed by Tukey's post hoc test or Kruskal-Wallis followed by Dunn's post hoc test. In (E) Two-way ANOVA followed by Tukey's post hoc test. *P < 0.05, **P < 0.01, ***P < 0.001, ****P < 0.0001. #P < 0.05 Student´s t-test (C).
